# Allosteric Regulation of a Synaptic Vesicle Glutamate Transporter

**DOI:** 10.1101/2022.07.26.501550

**Authors:** Fei Li, Jacob Eriksen, Juan A. Oses-Prieto, Yessica K. Gomez, Hongfei Xu, Janet Finer-Moore, Phuong Nguyen, Alisa Bowen, Andrew Nelson, Alma Burlingame, Michael Grabe, Robert M. Stroud, Robert H. Edwards

**Author notes:** Send correspondence to: R.H. Edwards at, R. M. Stroud at and F. Li at. These authors contributed equally (Fei Li, Jacob Eriksen).

## Abstract

Concentration of neurotransmitter inside synaptic vesicles (SVs) underlies the quantal nature of synaptic transmission. In contrast to many transporters, SV uptake of the principal excitatory neurotransmitter glutamate is driven by membrane potential. To prevent nonquantal efflux of glutamate after SV exocytosis, the vesicular glutamate transporters (VGLUTs) are allosterically inhibited by the neutral pH of the synaptic cleft. We have now determined high-resolution structures of VGLUT2 with a cyclic analog of glutamate bound that defines the mechanism of substrate recognition, a positively charged cytoplasmic vestibule that electrostatically attracts the negatively charged substrate, and modification by palmitoylation that promotes retrieval of the transporter after exocytosis. The structure also incorporates an extensive, cytoplasmic network of electrostatic interactions that acts as a gate. Functional analysis shows how this cytoplasmic gate confers the allosteric requirement for lumenal H^+^ required to restrict VGLUT activity to SVs.

## Introduction

The storage of neurotransmitter inside synaptic vesicles (SVs) enables their release by regulated exocytosis, conferring the quantal mode of neurotransmission^1^. The amino acid glutamate serves as the principal excitatory neurotransmitter in most neural systems and is packaged into SVs by highly selective but low affinity vesicular glutamate transporters (VGLUTs)^2–5^. Mammals express three closely related VGLUT isoforms with similar transport activity but different expression patterns^6–8^. Loss of VGLUT expression eliminates glutamatergic transmission^8, 9^.

The VGLUTs belong to the <u>S</u>olute <u>C</u>arrier 17 (SLC17) family of secondary active transporters^10^. Most secondary active transporters couple the concentration of substrate to the flux of another molecule, in many cases an ion^11^. Indeed, the highly homologous lysosomal sialic acid transporter sialin (41% sequence identity to VGLUT2) and the *E. coli* D-galactonate transporter DgoT (23% sequence identity) use H^+^ symport to drive organic anion transport into the cytoplasm^12–14^ (Extended Data Fig. 1). In contrast, the VGLUTs are not coupled to H^+^ but rather driven by membrane potential (ΔΨ), and transport in the opposite direction, moving glutamate from the cytoplasm into SVs, against a pH gradient^3, 15^. Equilibrating the anionic substrate according to ΔΨ enables the VGLUTs to concentrate glutamate 10-20 fold. Indeed, SVs contain >100 mM glutamate^16^, a concentration similar to that predicted by coupling to ΔΨ^17^. Despite the concentration of glutamate inside SVs, the VGLUTs do not leak glutamate even after dissipation of ΔΨ^18, 19^ and the mechanism responsible remains unknown.

SV exocytosis exposes VGLUTs to the positive outside cell membrane potential, which has the same orientation as the SV membrane potential and should drive non-vesicular glutamate efflux from the neuron. To prevent the degradation of quantal neurotransmission, glutamate transport at the plasma membrane is prevented by the change in environment, in particular the pH, which is neutral in the synaptic cleft rather than acidic as inside SVs^20^. Glutamate uptake by SVs also requires the efflux of Cl^-^ trapped by endocytosis, and the VGLUTs exhibit an associated Cl^-^ conductance as well as allosteric regulation by Cl^-2–4, 20–28^. These properties take advantage of the rapidly changing ionic conditions associated with the SV cycle to enhance and to control glutamate flux by the VGLUTs but the mechanisms for allosteric regulation by H^+^ and Cl^-^ remain poorly understood.

## Results

### Substrate recognition

The high rates of release observed at many synapses require mechanisms to recapture the transporters into recycled SVs and rapidly refill SVs with high concentrations of neurotransmitter. Direct measurement has shown that refilling with glutamate can occur within 15s^26^. High presynaptic concentrations of glutamate facilitate this process and require an appropriately tuned transport affinity. At the same time, the VGLUTs must discriminate between glutamate and similar compounds that also accumulate in the nerve terminal such as aspartate^29^. Consistent with these requirements, the VGLUTs transport glutamate with high specificity but with low apparent affinity (*K*_m_ ∼1-3 mM). However the VGLUTs do not bind or transport closely related aspartate^2^. To understand the basis for this specificity, we used a cyclic analog of glutamate, L-*trans*-1-amino-1,3-dicarboxy cyclopentane (ACPD), which is a competitive inhibitor of the VGLUTs that is transported with higher apparent affinity (*K*_i_ ∼*K*_m_ ∼0.23 mM) than glutamate^30^. To stabilize a substrate-bound conformation, we used a mutant (R181Q/E191Q) that impairs vesicular glutamate transport after reconstitution (Extended Data Fig. 2). Since high concentrations of Cl^-^ are also non-competitive inhibitors of glutamate transport^25^, we used low Cl^-^ (1 mM). The structure of the R181Q/E191Q VGLUT2-ACPD complex was determined using a previously described Fab^31^ to 3.0 Å resolution (Fig. 1**a-c**; Extended Data Fig. 3, 4**a**).

**Figure 1.**
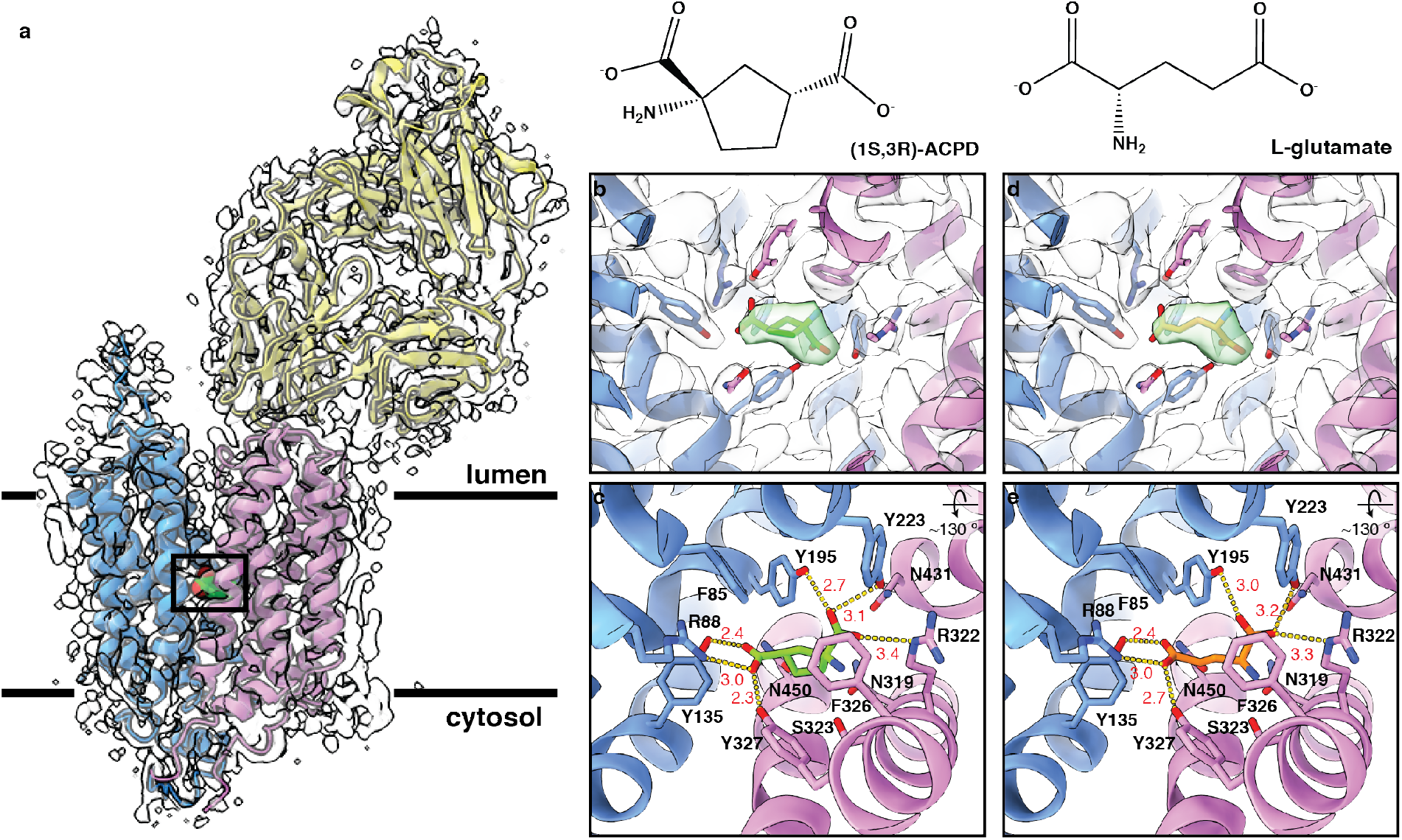
Substrate binding site. **a**, Overall structure of R184Q/E191Q VGLUT2-Fab complex bound to ACPD. N-domain is indicated in blue, C-domain in pink and Fab in yellow. ACPD is shown as a space-filling model in green. EM density is shown as white transparent surface. **b**,**d**, Density around the ligand binding site with ACPD (**b**) and glutamate (**d**) shown in the density (green). **c**,**e,** Magnified view of interactions between ACPD (**c**) and glutamate (**e**). Distances between interacting atoms are labeled in red. A cation-pi interaction between the amine group of ACPD and F326 is also observed with a distance of 3.3 Å.

The VGLUT2-ACPD complex adopts a lumen-open conformation with ACPD bound at the center of the protein between N- and C-domain helical bundles (Fig. 1**a-c**). The two carboxyl groups of ACPD are coordinated in almost symmetrical fashion, each by one arginine and two tyrosine residues. The 3’-carboxyl is coordinated by R88 and Y135 from the N-domain and Y327 from the C-domain, whereas the 1’-carboxyl is coordinated by R322 from the C-domain as well as Y195 and Y223 from the N-domain. Both domains thus coordinate glutamate to make multiple hydrogen bonds as well as electrostatic interactions with each carboxyl (Fig. 1**c**). Interestingly, the critical residue R322^25^ uses its *ε*-NH in a folded-back configuration, rather than its terminal *η*-NH_2_, to form an H-bonded salt bridge with the 1’-carboxyl (Fig. 1**c**). This configuration can rationalize the inability of VGLUTs to recognize aspartate^2^: with ∼3.4 Å between the 1’-carboxyl oxygen of ACPD and the *ε*-nitrogen of R322 (Fig. 1**c**), the shorter aspartate would not bridge the distance between R88 and R322. Considering the high similarity between glutamate and the cyclic analog ACPD, we replaced and refined glutamate into the density for ACPD (Fig. 1**d**), which preserved the same interactions with the extremities of glutamate (Fig. 1**e**). Previous studies showed that the amine group of glutamate is critical for recognition by the VGLUTs^2, 30^. The amine of ACPD lies 3.3 Å away from F326, forming a pi-cation interaction with the phenyl ring (Fig. 1**c**). In addition, an outer layer of polar residues including N431, N450 and S323 surrounds the recognition site and may also interact with the amino group of ACPD/glutamate in the fully occluded conformation.

As validation, ACPD was docked into the R184Q/E191Q structure using Schrodinger software. This resulted in a similar pose that preserves the interaction landscape with nearly identical ring and functional group placement (Extended Data Fig. 5**a**). We also docked glutamate into the wild-type apo structure (PDB ID 6V4D)^31^. The resulting top-ranked glutamate pose was again similar to the reported ACPD structure, with good alignment between the carboxyl groups but a different orientation of the amine group (Extended Data Fig. 5**b**). To probe the stability of these interactions, molecular dynamics (MD) simulations of glutamate with wild-type VGLUT2 and of ACPD with R184Q/E191Q VGLUT2 were carried out. The ACPD simulation was initiated from the structure and the glutamate simulation from the docked pose. Both MD simulations indicate that R88 and R322 form stable bidentate interactions with the ligand carboxyl groups (Extended Data Fig. 5**d, e, g**); as observed in the structure, the *ε*-NH and one *η*-NH_2_ of R322 form a salt bridge with the substrate. The amine group of ACPD engages F326 for the majority of the simulation but sometime flips and reorients towards N450 with a distance ∼4 Å (Extended Data Fig. 5**c-e**). Similarly, the amine of glutamate persistently contacts the sidechain oxygen of N450 and occasionally rotates to interact with F326 (Extended Data Fig. 5**f, g**), consistent with the importance of the amine group for recognition by VGLUT^2, 30^. MD simulation also indicates that glutamate remains highly flexible in the binding pocket and may reverse its conformation while preserving critical interactions with the flanking arginines; this flexibility may contribute to the high *K*_m_ of the VGLUTs.

### Substrate release

To activate postsynaptic receptors, SVs accumulate high millimolar concentrations of neurotransmitter. This requires discharge of the neurotransmitter into the lumen with high concentrations of neurotransmitters already as the SV fills up. To determine the structural changes associated with glutamate discharge by the VGLUTs, we compared the two VGLUT2 structures: R184Q/E191Q with the substrate analogue ACPD and WT without substrate (Fig. 2). To simplify the comparison, we determined the WT structure under the same low Cl^-^ condition to 3.3 Å resolution (Extended Data Figs. 6, 4**b**). Both structures open to the lumen and thus differ primarily in the presence or absence of substrate. In the ACPD structure, the lumenal half of the C-domain rotates slightly over ACPD (Fig. 2**a, b**). Substrate-binding residues from the N-domain do not change between the two structures whereas TM7 of the C-domain exhibits the largest conformational change. L330, Y327, and F326, bulky residues on the lumenal side of TM7, rotate significantly to enclose ACPD in the central binding site. In addition, S323, R322, and N319 on the cytoplasmic side of TM7 translate toward the N-domain (Fig. 2**b**).

**Figure 2.**
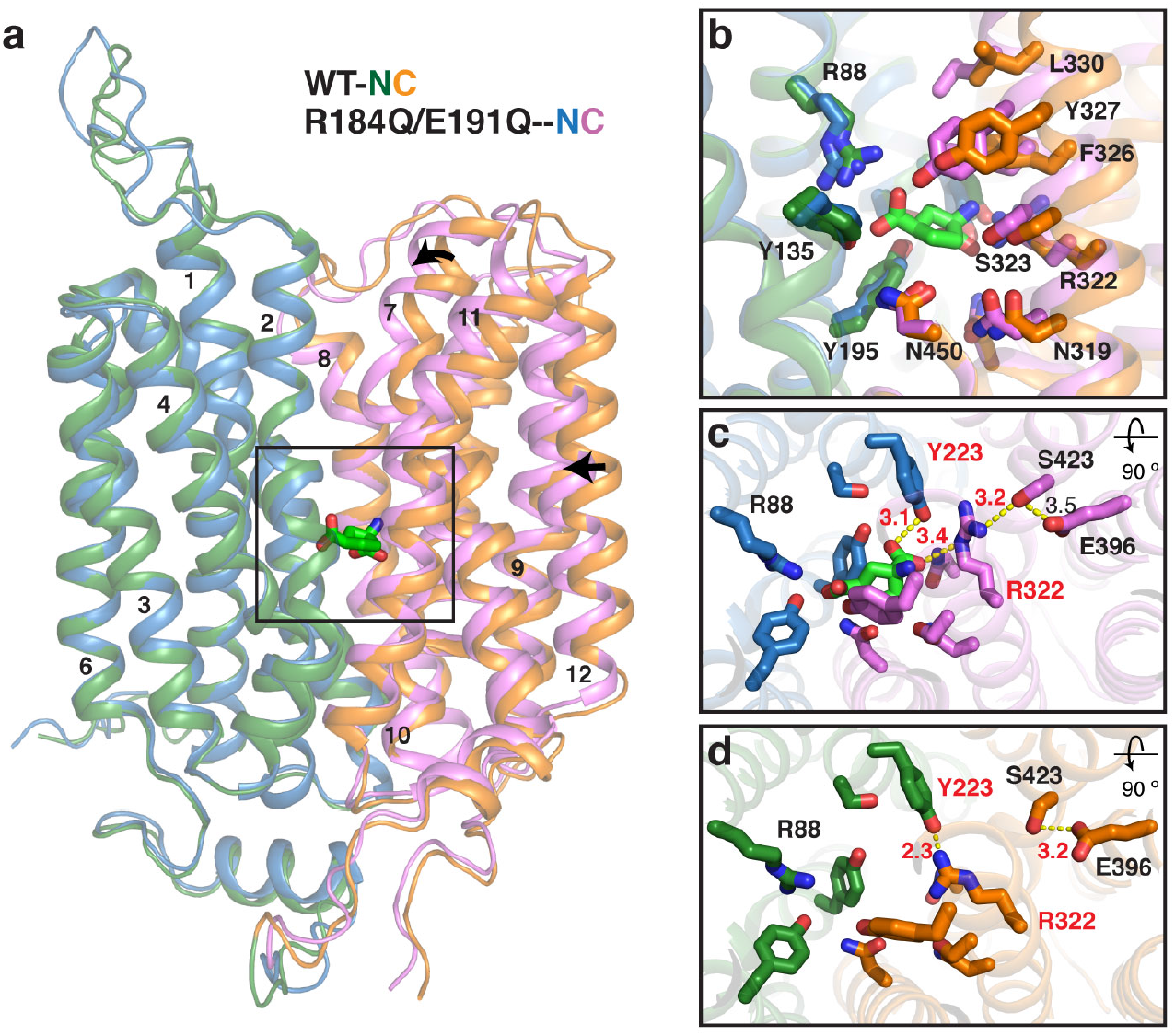
Mechanism for substrate release. **a**, Structure of R184Q/E191Q VGLUT2 adopts a similar but more closed lumen-open conformation than WT, with the lumenal section of TM7 exhibiting the largest change. **b**, Bulky side chains on TM7 (Y327, F326, L330) play the major role in closing the substrate binding site to the lumen. **c**, Magnified view of the substrate-binding site in R184Q/E191Q VGLUT2 with ACPD bound. **d**, Substrate binding site of the apo WT VGLUT2 structure.

Differences between the two structures in the interaction network around the essential R322 indicate a mechanism that accompanies substrate release into the vesicle lumen (Fig. 2**c, d**). In the ACPD-bound structure, both Y223 and R322 coordinate the substrate carboxyl whereas the *η*-NH_2_ of R322 hydrogen bonds with S423 (Fig. 2**c**). In the apo structure, by contrast, S423 is sequestered by E396, releasing R322 to hydrogen bond with Y223 (Fig. 2**d**). In this configuration, the substrate 1’-carboxyl group loses two of three critical interactions with the transporter, reducing the enthalpic component and favoring discharge into the lumen.

### An asymmetric transport pathway

In the absence of neural activity, SVs exhibit very low rates of glutamate leakage even with dissipation of ΔΨ, the driving force for glutamate uptake^18, 19^. The structures of VGLUT2 reveal an asymmetry in the pathways leading to the central substrate binding site that may reduce leakage. On the lumenal side, residues lining the central cavity are significantly more hydrophobic than those on the cytoplasmic side, especially along TM7 of the C-domain (Fig. 3**a**). Taking advantage of the cytoplasm-open structure of DgoT^14^, we generated models of the cytoplasm-open conformation for both R184Q/E191Q and WT VGLUT2. In both models, the N- and C-domains of VGLUT2 adopt structures very similar to their respective domains in DgoT, with conserved residues at identical positions (Extended Data Fig. 7**a, b**). However, the surface electrostatics of VGLUT2 shows strikingly different paths to the central cavity between cytoplasm- and lumen-open conformations. In the cytoplasm-open conformation, a continuous positively charged path (Fig. 3**d**) electrostatically attracts entry of anionic glutamate from the cytoplasm. In the lumen-open conformation, a series of acidic residues (D97, E122, E338, E339, and E344) line the mouth of the lumenal cavity disfavoring an anion in the vestibule, and the path to the central cavity is much more hydrophobic (Fig. 3**a, b, c**). In general, residues lining both cavities show low sequence conservation across the SLC17 family, however the vesicular nucleotide transporter (VNUT), also driven by ΔΨ, has three acidic residues on the lumenal surface of TM7 similar to E338 and E339 in VGLUT2 and the homologous residues in VGLUT1 and 3 (Fig, 3**a**, Extended Data Fig. 1). In contrast, the H^+^ coupled symporter DgoT that drives substrate into the cytoplasm has a more symmetric binding cavity with positive charge extending from the central recognition site to both sides of the membrane (Extended Data Fig. 7**c-e**). The negative charge and hydrophobicity of the VGLUTs impose a barrier to glutamate rebinding from the lumen, providing a mechanism for the low rates of leakage. This barrier may also impede glutamate release into the vesicle lumen, requiring other mechanisms for release such as the change in conformation of R322 described above.

**Figure 3.**
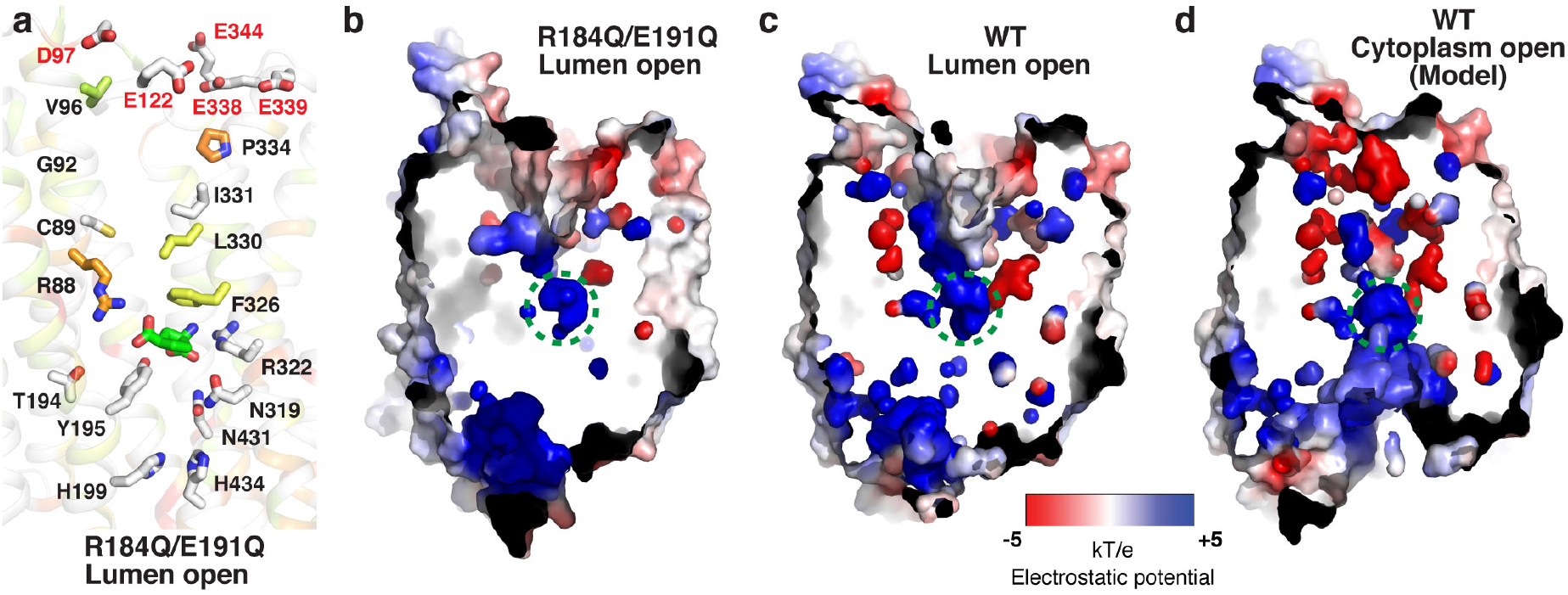
Asymmetrical paths to the central cavity facilitate directional transport driven by ΔΨ. **a**, Structure of the binding cavity R184Q/E191Q VGLUT2 colored with sequence conservation as in Extended Data Fig. 1 (red the most conserved and no color the least conserved). **b**, Electrostatic surface of the lumen-open R184Q/E191Q structure without ACPD. **c**, Electrostatic surface of the apo WT structure in lumen-open conformation. **d**, Electrostatic surface of a model for WT in the cytoplasm-open conformation. Locations of the substrate binding site in (**b**-**d**) are outlined by a dashed green circle.

### Palmitoylation regulates VGLUT trafficking

The high-quality map of R184Q/E191Q (Extended Data Fig. 4**a**) identifies a lipid density at conserved C62 of the cytoplasmic N-terminus (Fig. 4**a, b**). A palmitate molecule is well-defined in the density, and another weaker density is also observed at adjacent residue C64 (Fig. 4**b**). Palmitoylation is a lipid modification observed on integral as well as peripheral membrane proteins that can regulate their association with membranes or membrane microdomains^32^. Palmitoylated proteins include the presynaptic cysteine string protein, t-SNAREs syntaxin, SNAP25 and v-SNARE synaptobrevin, but the role of palmitoylation in the exo- and endocytic recycling of SV membrane proteins remains poorly understood^33^.

**Figure 4.**
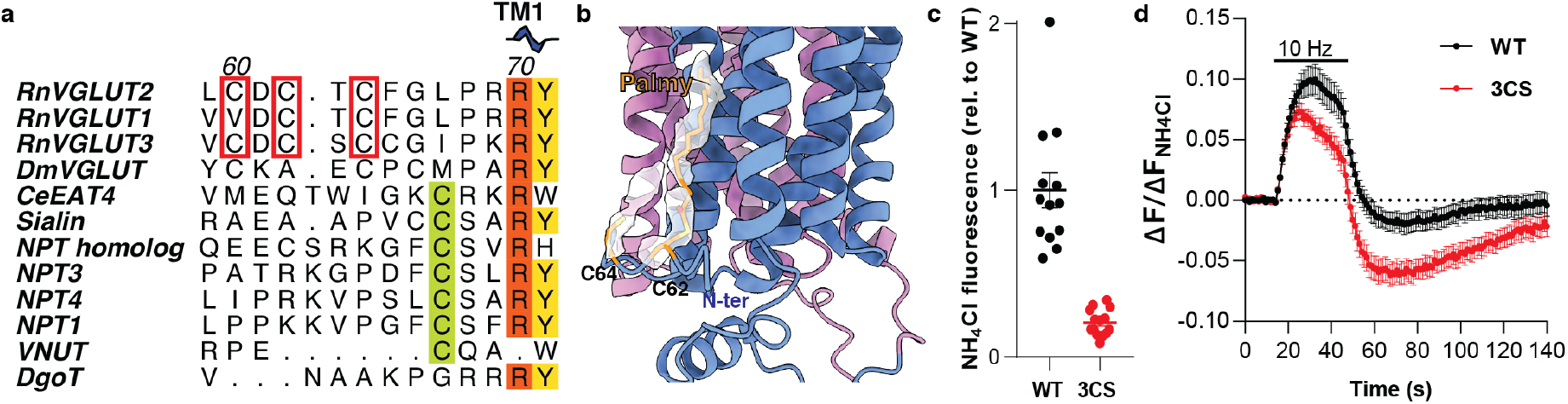
VGLUT2 is palmitoylated at C60, C62 and C64. **a**, Alignment of SLC17 protein cytosolic N-termini proximal to TM1, which contains the palmitoylated cysteine residues (red boxes). C62 and C64 are conserved in all three mammalian VGLUTs whereas C60 is conserved in VGLUT2 and 3 but replaced by a valine in VGLUT1. Other SLC17 family members have a conserved cysteine closer to TM1 (green). Sequence alignment is colored to indicate conservation as in Extended Data Fig. 1 (red most conserved and no color least conserved). **b**, Structure of R184Q/E191Q VGLUT2 with the C62 and C64 palmitoylate densities. N-domain is blue and C-domain pink. Two lipid densities (white) are resolved on C62 and C64. A palmitoyl group (palmy) fits into the density at C62. **c**, Total pHluorin fluorescence (in NH_4_Cl) from hippocampal neurons (DIV 13-15) expressing WT or 3CS mutant (C60S, C62S and C64S) VGLUT2-pHluorin. **d**, Normalized pHluorin response to 10 Hz stimulation for 30 s. (**c**,**d**; n=13-14 coverslips per condition).

To confirm the identity of lipid modification and the sites modified, we analyzed the purified VGLUT2 used for cryo-EM by mass spectrometry. We found modification of three N-terminal cysteines (C60, C62 and C64) by palmitic acid (Extended Data Fig. 8), consistent with putative sites identified from mouse brain^34^. To determine the role of palmitoylation, we monitored exo- and endocytosis of VGLUT2 in cultured neurons by live cell imaging of a pH-sensitive form of GFP (pHluorin) fused to the lumenal surface of the transporter. Fluorescence of the pHluorin is quenched by the low pH inside SVs and increases with exposure to the higher external pH during exocytosis^35, 36^. Mutation of all three cysteines to serine (3CS) drastically reduces the expression of VGLUT2 (Fig. 4**c**). Corrected for reduced expression, the mutations do not impair the initial exocytic response of VGLUT2 to stimulation (Fig. 4**d**). However, the mutations result in a greater fluorescence decline after stimulation (Fig. 4**d**). This decline reflects both the re-acidification that accompanies endocytosis of SV proteins and their dispersion into the axonal plasma membrane. In the case of the 3CS mutant, the subsequent return back up to baseline takes longer than for wild-type, suggesting increased axonal dispersion that slows endocytic recycling, reclustering and the regeneration of SVs. Thus, palmitoylation serves to anchor VGLUT2 near the site of exocytosis and promote the efficient retrieval required for high rates of glutamate release.

### The cytoplasmic gate and regulation by lumenal pH

The alternating access mechanism of transport generally involves the sequential opening and closure of gates on each side of the membrane. The high-quality density map for R184Q/E191Q VGLUT2 now allows us to unambiguously resolve solvent-exposed flexible residues, revealing an extensive ionic interaction network that appears to act as the cytoplasmic gate. This network comprises alternating positively charged (arginine) and negatively charged (glutamate or aspartate) residues that interact with each other and span both N- and C-domains as well as intracellular helix 1 (ICH1) located between TM6 and 7 (Fig. 5**a, c**). E278 in ICH1 binds R70 in TM1, which also interacts with E210 at the beginning of TM5. All three residues are highly conserved through the entire SLC17 family (Fig. 5**b**). Consistent with their importance, mutation of the residue equivalent to VGLUT2 R70 in sialin (R39C) causes Salla disease by impairing lysosomal efflux of sialic acid^12, 13^. In VGLUT2, the network continues with interaction of E210 and R213 at the beginning of TM5 and of R213 with D371 at the bottom of TM8, an interaction that bridges the N- and C-domains (Fig. 5**c**). D371 is in turn surrounded by three arginines, R374, and R385 from the C-domain as well as R213. Similarly, R374 and R385 from the C-domain and R211 from the N-domain surround D436 from the C-domain. These extensive ionic interactions stabilize the closed cytoplasmic gate in this lumen-open conformation. D371 is part of MFS cytosolic motif A [Gxxx(D/E)(R/K)xGx(R/K)(R/K)], which has been shown to stabilize the outward-open conformation and promote transport activity^37, 38^.

**Figure 5.**
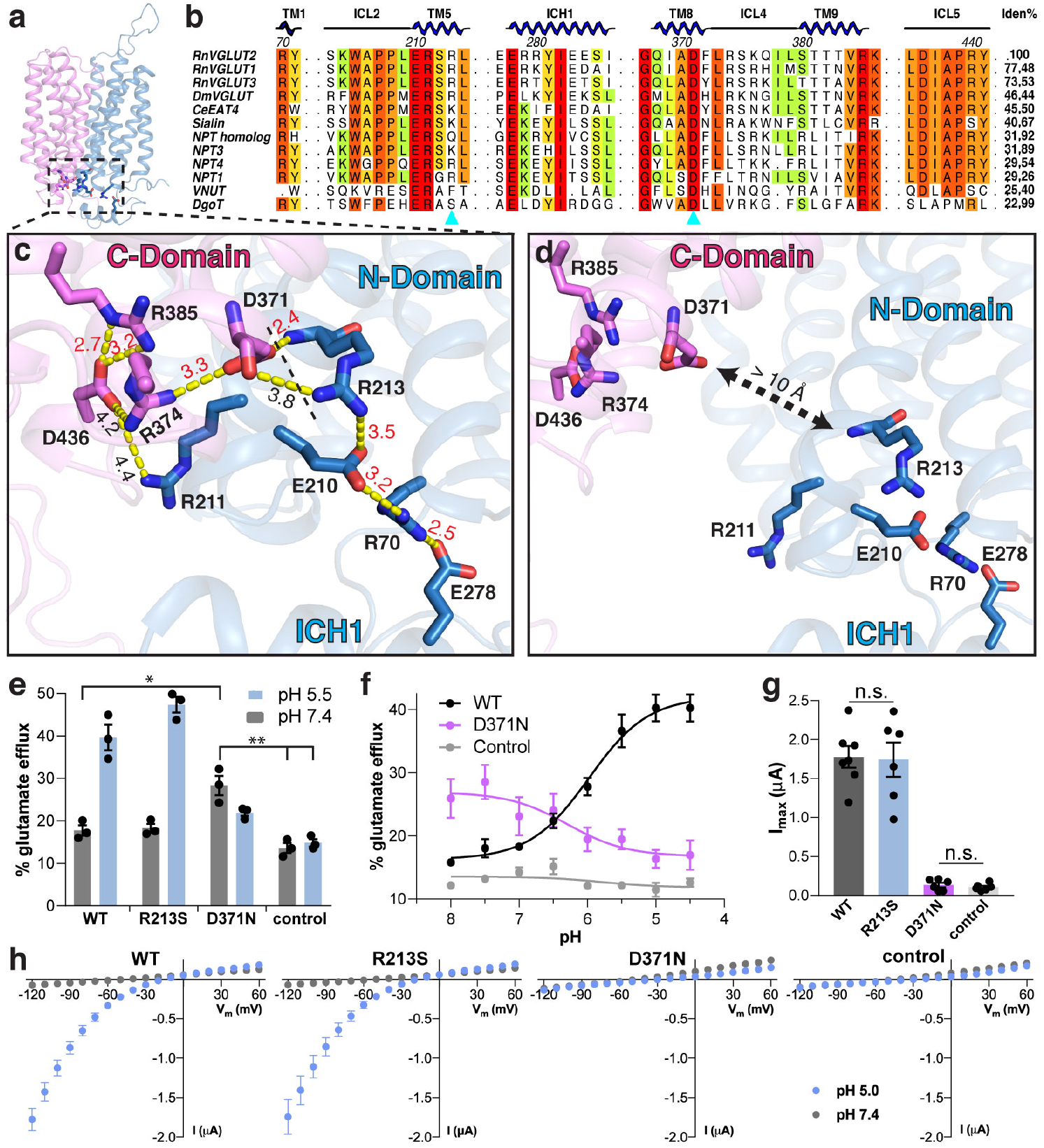
Cytoplasmic ionic interaction network. **a**, Overall structure of R184Q/E191Q VGLUT2 with the cytoplasmic interaction network (box) magnified in **c**. N-domain is blue and C-domain pink. **b**, Sequence alignment of the cytoplasmic interaction network in VGLUT homologs and other SLC17 family members. Color indicates conservation, with red the most conserved and no color the least. R213 and D371 (cyan triangles) interact across N- and C-domains. **c**, Ionic interactions between N- and C-domains as well as the ICH1 helix on the cytoplasmic face of R184Q/E191Q VGLUT2 in the lumen-open conformation. Distances between interacting atoms are labeled in red and between potential interactions in black. **d**, In the cytoplasm-open conformation of VGLUT2 (modeled on DgoT), the interaction between D371 and R213 is broken. **e**, VGLUT2 activity measured as ^3^H-glutamate efflux from HEK293T cells expressing WT, R213S and D371N plasma membrane-targeted (pm) VGLUT2. R213S transport activity resembles WT but D371N shows reduced activity at pH 5.5 and increased activity at pH 7.4. **f**, pH response curve for glutamate efflux from cells expressing WT, D371N pmVGLUT2 and untransfected controls. WT shows potentiation by protons (EC_50_: pH = 6.0 ± 0.025) whereas D371N shows inhibition by protons (IC_50_: pH = 6.2 ± 0.27). n=3 for **e** and **f**. **g**, Maximum inward currents (at −120 mV) from oocytes expressing WT, R213S, D371N pmVGLUT2-HA and uninjected controls (n=6 oocytes per condition). **h**, Individual I-V curves for each of the conditions in **g**. n.s. p > 0.05, ∗ p < 0.05; ∗∗ p < 0.01, two-way ANOVA with correction for multiple comparisons.

To determine the functional significance of the cytoplasmic interaction network, we neutralized the two interacting residues (R213 and D371) that bridge N- and C-domains by mutation. The backbone amine group of R213 interacts with the sidechain carboxyl of D371, and the sidechain of R213 may form a weaker interaction with the backbone carbonyl of D371 (Fig. 5**c**). We assessed the effect on glutamate transport using an assay for efflux from HEK293t cells expressing a plasma membrane-targeted version of VGLUT2^39^. In this assay, VGLUT-mediated efflux of preloaded ^3^H-glutamate depends on low external pH that mimics the lumenal pH of SVs. The R213S mutation does not affect glutamate transport, with activity similar to WT (Fig. 5**e**), consistent with an interaction that involves the backbone of R213. However, D371N both reduces glutamate efflux and changes its pH dependence. In contrast to WT, D371N VGLUT2 shows more activity at high rather than low external pH (Fig. 5**e**). Thus, D371N reverses the pH dependence of the VGLUTs. WT VGLUT2 shows activation by protons with a sigmoidal dose response (EC_50_; pH = 6.0 ± 0.025) whereas D371N shows inhibition by protons but with a similar pH sensitivity (IC_50_; pH 6.2 ± 0.27) (Fig. 5**f**). The differential effect of R213S and D371N is consistent with a critical role for the interaction between the backbone of R213 and the sidechain of D371. Since the VGLUTs exhibit an associated Cl^-^ conductance that is not coupled to glutamate flux, we also tested the effect of the mutations on this property using oocytes expressing plasma membrane-targeted VGLUT2^20^. The R213S mutation has no effect on these currents but D371N eliminates them (Fig. 5**g, h**; Extended Data Fig. 9). Thus, the integrity of the cytoplasmic gate both influences the allosteric regulation by lumenal H^+^ and enables the Cl^-^ conductance associated with VGLUTs.

## Discussion

The structures reported here indicate how the VGLUTs recognize glutamate with the low apparent affinity but high specificity required for the rapid refilling of SVs. One arginine and two tyrosine residues contact each of the carboxyl groups and the distance between these two sites dictates the length requirement for substrate recognition that excludes a role for aspartate as neurotransmitter^29^. The arginine in TM1 is conserved through many members of the SLC17 family, consistent with a role in anion transport (Extended Data Fig. 1). However, the arginine in TM7 occurs only in the VGLUTs and not in any other members of SLC17, which transport substrates such as urate, p-aminohippurate and sialic acid that carry a single negative charge. The amine in glutamate forms either a cation-pi interaction with F326 or a hydrogen bond with N450 and flipping between these two orientations observed by MD simulation is consistent with the low apparent affinity of the VGLUTs.

Comparison of the two VGLUT2 structures with and without substrate suggests a mechanism that accompanies substrate discharge into the SV lumen. In the apo structure, two of the three residues recognizing the 1’ carboxyl of ACPD interact with each other rather than with substrate. This change may result from the loss of substrate, or it may promote substrate discharge. A mechanism to promote substrate release may become increasingly important as the SV fills with glutamate. In addition, after glutamate discharge into the lumen, Y223 may serve to neutralize R322 to reduce electrostatic repulsion, thus promoting reorientation of the unloaded carrier to the cytoplasm. Similarly, substrate glutamate serves to neutralize the positive charges on R88 and R322, enabling translocation of the loaded carrier from the cytoplasm back to the lumen and completing a productive transport cycle. Neutralization of R88 and R322 may thus play an important role in reorientation of both loaded and unloaded VGLUT.

An asymmetry in the main cavity of VGLUT2 suggests a mechanism to regulate the rates of substrate rebinding in the lumen. As shown in the model based on DgoT, the main cavity of cytoplasmically oriented VGLUT2 contains a positively charged path to the substrate recognition site which should promote anion entry. In the lumenal orientation, however, the main cavity is lined with hydrophobic residues. The hydrophobicity will slow rebinding from the lumen and, along with the requirement for allosteric activation by H^+^, may contribute to the low rates of leakage observed after inhibition of the v-type H^+^-ATPase^18, 19^. The hydrophobicity may also slow substrate discharge into the lumen. but this could be overcome by other mechanisms such as the rearrangement of substrate-recognizing residues described above.

The structure also reveals palmitoylation of the VGLUT2 N-terminus. Many SV proteins undergo palmitoylation but the significance has remained unclear^33^. We now find that this modification serves to restrict the movement of VGLUT2 in the plasma membrane after exocytosis, thus promoting its rapid retrieval by endocytosis and the regeneration of SVs capable of glutamate storage. Presumably, palmitoylation restricts VGLUT2 to a membrane microdomain^32^ surrounding the release site, predicting a similar role in the endocytic recapture of other SV proteins.

Based on the structures, we propose the following mechanism for ΔΨ-driven transport by the VGLUTs (Fig. 6). In the lumen/outward-open conformation, the VGLUTs are stabilized by extensive ionic interactions across N- and C-domains at the cytoplasmic face of the protein. Neutralization of negative charges on the lumenal side of the VGLUT by H^+^ binding enables closure of a putative lumenal gate, rupturing the cytoplasmic network and allowing reorientation toward the cytoplasm. Glutamate then enters the substrate binding site through the positively charged, cytoplasmically oriented main cavity. Transition through an occluded state that involves recognition of each glutamate carboxyl by both N- and C-domains then reorients the transporter to the SV lumen, re-establishing the cytoplasmic network. In this conformation, a change in the interaction with R322 disrupts glutamate recognition, allowing substrate to move down its electrochemical gradient into the SV lumen. The negative charge and hydrophobicity of the lumenal vestibule then slow glutamate rebinding from loaded SVs even if ΔΨ decreases. Empty transporter then returns to the cytoplasm-open conformation with R322 bound to Y223.

**Figure 6.**
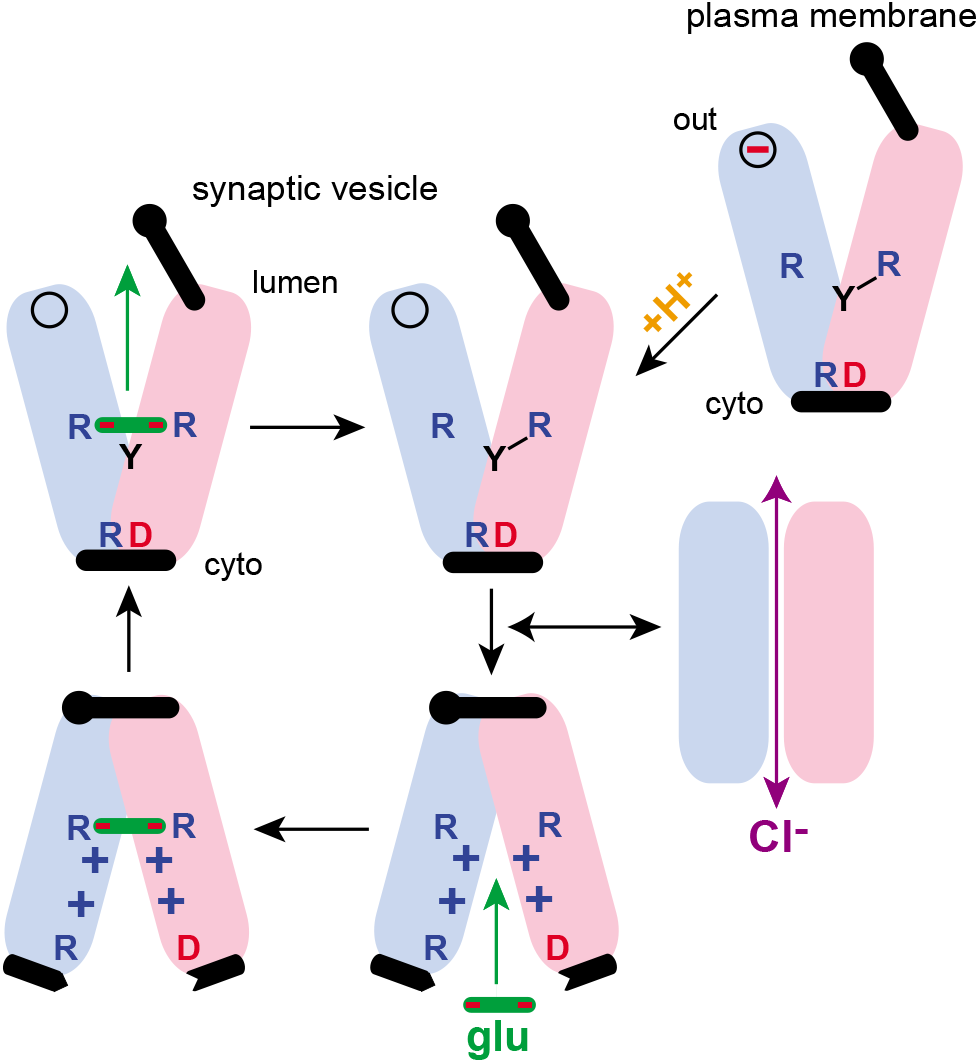
Proposed transport cycle for VGLUT2. At neutral pH in the plasma membrane, VGLUT is stabilized in the lumen-open conformation with cytoplasmic gate closed (upper right panel). Protonation of a lumenal site (indicated here with a minus inside a circle) at the low pH of SVs enables closure of the lumenal gate, disrupting the interaction between R213 and D371, rupturing the cytoplasmic gate and driving reorientation to the cytoplasm (lower middle panel). In this conformation, the anionic substrate glutamate enters the binding site through the positively charged cavity (lower panels) and binds to R88 and R322, driving reorientation back to the lumen-open conformation (upper left panel). Rearrangement of R322 to interact with Y223 accompanies glutamate release into the SV lumen. The balance between lumenal and cytoplasmic gates controls conformational dynamics and hence transport activity. In WT VGLUT, the cytoplasmic gate stabilizes the lumen-open conformation. In this case, lumenal protonation lowers the energy barrier for closure of the lumenal gate, enabling reorientation to the cytoplasm and alternating access. However, weakening of the cytoplasmic gate (e.g., by the D371N mutation) shifts the equilibrium to favor the cytoplasm-open conformation, and reorientation to the lumen becomes rate-limiting. In this case, lumenal deprotonation favors opening of the lumenal gate, allowing completion of the transport cycle. The cytoplasmic gate thus tunes allosteric regulation from the lumen, enabling pH to coordinate glutamate transport with the SV cycle. In addition, weakening of the gate by D371N eliminates the VGLUT-associated Cl^-^ conductance, suggesting that the mutation destabilizes a transient intermediate required for the conductance.

Since their identification, the VGLUTs have been recognized to undergo potent allosteric regulation^2, 22, 26^, but the mechanisms have remained unclear. Lumenal protons activate both glutamate transport and the associated Cl^-^ conductance^20, 39^, restricting the activity of the VGLUTs to SVs and preventing nonquantal release. Our data suggest a mechanism for allosteric activation by lumenal pH that depends on tuning of the strength of lumenal and cytoplasmic gates. The D371N mutation weakens the cytoplasmic gate between N- and C-domains and reverses the lumenal pH dependence of glutamate transport by VGLUT2. The midpoint of pH activation does not differ from WT, suggesting involvement of the same titratable residue. Lumenal H^+^ binding inhibits transport by the D371N mutant but activates WT VGLUT2.

How can a cytoplasmic mutation affect the role of lumenal protonation? At lumenal pH ∼7, cytoplasmic gate closure stabilizes the outward open orientation of WT VGLUTs, and we hypothesize that protonation promotes closure of a lumenal gate required for reorientation to the cytoplasm. D371N weakens the cytoplasmic gate, thereby promoting closure of the lumenal gate and eliminating the requirement for activation by lumenal H^+^. Indeed, activation of D371N at higher pH suggests that deprotonation weakens the lumenal gate, thereby restoring the balance of strength between the two gates required for alternating access. Thus, D371N and WT VGLUT2 both undergo protonation in the lumen, but the strength of the cytoplasmic gate determines the overall effect on transport. In the bacterial permease lacY, the mutation of periplasmic residues rescues cytoplasmic motif A mutations^37^, supporting a role for the balance between opposing gates, although not in the context of regulation. The allosteric regulation of VGLUTs by Cl^-^ may involve a related mechanism. Chloride binds to a similar cytoplasmic interaction network in the cytoplasm-open conformation of two other MFS transporters (the glucose transporter GLUT1 and the plant sugar transporter STP10) but not in their outward-open conformation with cytoplasmic gates closed^40, 41^. In the VGLUTs, cytoplasmic Cl^-^ may thus be required to weaken the cytoplasmic gate and hence promote the cytoplasmic open orientation required for substrate binding. The associated Cl^-^ conductance was abolished by the D371N mutation, showing that it requires a conformation disfavored by weakening of the cytosolic gate. The two functions of the VGLUTs are thus separable: the alternating access required for glutamate transport depends on the appropriate balance between cytoplasmic and lumenal gates whereas the Cl^-^ conductance involves a specific intermediate, indicating the potential for differential regulation of these two activities in the course of SV filling. The insight into mechanism provided by structure will enable future investigation of the role for these properties in neurotransmission.

## Acknowledgements

We thank Dr. David Bulkley, Dr. Zanlin Yu, Mr. Glenn Gilbert and Mr. Matt Harrington at the UCSF cryo-EM facility and Dr. Corey Hecksel, Dr. Patrick Mitchell, Dr. Lydia-Marie Joubert and Ms. Lisa Dunn at Stanford-SLAC Cryo-EM Center (S^2^C^2^) for their support in data acquisition and computation. We thank Dr. Daniel Cawley at VGTI (OHSU) for generating the mAb and advice on working with the antibodies. This work was supported by R01NS089713 to R.M.S and R.H.E. and R37MH50712 to R.H.E. F.L was supported by postdoctoral fellowships from American heart Association (17POST33660928) and NIMH (K99MH119591). The UCSF EM facility was supported by NIH grants S10OD020054 and S10OD021741. Some of this work was performed at the Stanford-SLAC Cryo-EM Center (S^2^C^2^), which is supported by the National Institutes of Health Common Fund Transformative High-Resolution Cryo-Electron Microscopy program (U24 GM129541). The content is solely the responsibility of the authors and does not necessarily represent the official views of the National Institutes of Health. Structural biology applications used in this project were compiled and configured by SBGrid^42^. Mass Spectrometry was provided by the Mass Spectrometry Resource at UCSF (A.L. Burlingame, Director) supported by the Dr. Miriam and Sheldon G. Adelson Medical Research Foundation (AMRF) and the UCSF Program for Breakthrough Biomedical Research (PBBR).

## Author Contributions

F.L, J.E, R.M.S and R.H.E conceived the project. P.N, A.N, A.B, J.E and F.L expressed the protein. F.L purified the protein, prepared all samples for EM, acquired cryo-EM data at UCSF and SLAC cryo-EM facilities and determined the structures. J.E, F.L and H.X. performed the functional experiments. J.A.O and A.B performed the mass spectrometry experiments. Y.K.G and M.G performed the docking and molecular dynamic simulation. F.L, J.F.M and R.M.S analyzed the structure. F.L, R.H.E, and R.M.S wrote the manuscript with input from all authors.

## Online Methods

### Protein expression and purification

WT and R184Q/E191Q VGLUT2 were expressed in *Spodoptera frugiperda* (*Sf9*) cells (Expression System, #94-0015). High-titer recombinant baculovirus (>10^9^ viral particles/mL) was obtained using the Bac-to-Bac Baculovirus Expression System (Thermo Fisher Scientific^®^) as previously described^31^. *Sf9* cells at a cell density of (2–3) ×10^6^ cells/mL were infected with P_1_ virus at a multiplicity of infection of 3. Cells were harvested by centrifugation 48 h after infection and stored at −80 °C until use. The frozen cell pellets were gently sonicated in lysis buffer containing 50 mM Tris (pH 7.4), 400 mM NaCl, 5 mM dithiothreitol (DTT), 1 mM phenylmethylsulfonyl fluoride (PMSF), DNase, and EDTA-free protease inhibitor cocktail tablets (Sigma^®^). 5 M NaCl stock solution was added to the broken cells to a final concentration of 1 M and the cells were washed by repeated douncing with a dounce homogenizer. Membranes were then sedimented at 200,000*g* and resuspended in buffer A (50 mM Tris, 400 mM NaCl, 10% glycerol). Subsequently, 1% (w/v) *n*-dodecyl-*β*-d-maltopyranoside (DDM, Anatrace^®^) and 0.2% (w/v) cholesteryl hemisuccinate (CHS, Sigma^®^) were added to solubilize the membranes at a detergent/protein ratio of 0.22 (w/w) by stirring at 4 °C for 4 h. WT protein with a 10x his tag at the C-terminus was purified as previously described^31^. R184Q/E191Q VGLUT2 with a twin-strep tag at the N-terminus was purified as previously described for reconstitution^20^ except DDM/CHS was used for solubilization and purification at the same concentrations as for WT VGLUT2. Purified proteins were concentrated with a 100 kDa Amicon concentrator (Millipore^®^) to 1-2 mg/ml and frozen at −80 °C in aliquots of ∼25-50 μL until used to form a complex with the Fab. VGLUT2 proteins were incubated with ligand in buffer for 30 mins before adding the Fab (8E11 or 9C6) to avoid interference of the Fab with conformational change. Purified Fab was then added to the mixture and incubated on ice for another 30 min before purifying the complex on a Superdex 200 Increase column (GE healthcare^®^) in size exclusion chromatography (SEC) buffer (20 mM HEPES, 150 mM NaCl, pH 7.4 adjusted with NaOH and 0.01% GDN) with additional ligand. For the R184Q/E191Q-ACPD complex, 30 mM ACPD in SEC buffer was used for the incubation and 1 mM ACPD was added to the SEC running buffer. For WT VGLUT2, 2.77 mM clodronate^43^ was used during incubation and 263 µM clodronate added to the SEC running buffer. However, no additional density for clodronate was observed in the structure. We therefore denote the structure as WT-apo.

### Cryo-EM sample preparation and data acquisition

Freshly purified VGLUT2-Fab complex (3 μL) was concentrated to 0.5 – 1 mg/ml and applied to Quantifoil^®^ holey carbon grids glow-discharged with an EMS 700 glow discharge device (UCSF EM core) using mixed air at 10-20 mA for 30 seconds. Copper 1.2/1.3-400 mesh grids were blotted with grade 595 standard Vitrobot filter paper for 4-7 s at 10 °C and 100% humidity using a Vitrobot^®^ Mark IV, followed by rapid plunging into liquid ethane cooled by liquid nitrogen. The dataset for R184Q/E191Q-ACPD was collected at UCSF and the dataset for WT VGLUT2 was collected through S^2^C^2^ at SLAC with the parameters shown in Table S1.

### Image processing

Micrographs were corrected for beam-induced drift using MotionCor2^44^. The contrast transfer function (CTF) parameters for each micrograph were determined using CTFFIND4^45^ and micrographs of poor quality (CTF lower than 6 Å) were discarded. All subsequent processing was carried out in RELION 3.0^46^ and cryoSPARC^47^. Similar strategies were used for WT and R184Q/E191Q VGLUT2 data as illustrated in Extended Data Fig. 3 and 6. Briefly, particles were picked from good micrographs using templates previously generated. 2-3 rounds of 2D classification with 4x binned particles were used to remove junk particles. The particle stack was then re-extracted as 2x binned particles and aligned to a single class in RELION. 6 parallel 3D classifications were then initialized from parameters at iteration 20-25 from the previous 3D classification. For the R184Q/E191Q dataset, good, unique particles from each run were selected, combined and subjected to another round of multi-reference 3D classification. For the WT dataset, 2 classes that resulted in the highest resolution reconstructions were selected and imported into cryoSPARC. Each class was further classified into 2 classes using the *ab initio* reconstruction scheme. Particles classes with high resolution features were then selected and refined in RELION using gold-standard 3D auto-refine and Bayesian polishing strategy^48^. Polished particles were then auto-refined in RELION with a soft mask applied to the protein region including VGLUT2 and the Fab. The same polished particle stack was also imported into CryoSPARC and refined in parallel using the non-uniform refinement strategy. Final maps obtained in both RELION and CryoSPARC achieved similar resolution and quality with minor variations in the flexible loop 1 region. Maps generated in RELION generally resolved more density in this region as it is included in the manually generated mask. Statistics of the final maps generated were calculated using the Fourier shell correlation (FSC) criterion and a threshold of 0.143 using Phenix_comprehensive_validation tool^49^ and reported in Table S1.

### Model building and structure refinement

Final maps generated by RELION were sharpened with Phenix Auto-sharpening before used for model building and refinement in Phenix. Sharpened maps generated in cryoSPARC were used directly. The structure of WT VGLUT2 in high Cl^-^ (150 mM) (PDB# 6V4D) was used as the initial model for building in Coot^50^. The local areas of the built models were first refined within Coot using real-space refinement with Ramachandran and geometry constraints. The full models of WT and R184Q/E191Q VGLUT2 were then refined using Phenix.real_space_refine without imposed constraints^49^. The density corresponding to loop 1 between TM1 and TM2 is of much lower quality. As a result, only the main chain was built in this area. Figures were prepared using Pymol and Chimera^51^.

### Docking

The Small Molecule Drug Discovery Suite 2017-4 (Schrödinger LLC) was used for all docking calculations. For docking to the wild-type protein, the previously published high Cl^-^ structure (PDB# 6V4D) was used alongside the R184Q/E191Q structure reported here. Both proteins were prepared for docking, with protonation states assigned at pH 7.4 ± 0.1 and missing side chains and loops modeled using the Protein Preparation Wizard workflow. ACPD coordinates were extracted from the R184Q/E191Q map and saved as a separate PDB, then protonation state and tautomers were assigned at pH 7.4 ± 0.1 using Epik in Maestro LigPrep with the OPLS3 forcefield. Glutamate was drawn with Maestro v11.4, followed by identical processing with Maestro LigPrep. For docking ACPD into R184Q/E191Q with a flexible mode, a 20×20×20 Å^3^ docking grid was centered around the putative binding site. The flexible mode allowed bond rotation and ring conformation sampling while penalizing non-planar amide groups. To dock glutamate into WT VGLUT2, a docking grid was centered around the same putative binding site. In all instances, the protein was held rigid while the Glide XP scoring function was employed and a maximum of 10 optimal poses were returned. Representative poses were picked based on a combination of visual inspection, chemical intuition and score.

### Molecular dynamics simulations

Two systems were used to initiate simulations: one with R184Q/E191Q VGLUT2 and ACPD, and another with wild-type VGLUT2 and glutamate. Initial atomic coordinates of the glutamate and WT protein were taken from the Glide docking outputs, whereas the R184Q/E191Q VGLUT2 and ACPD coordinates used directly from the cryo-EM structure. The CHARMM-GUI bilayer builder module was used to place each protein in a 1:1 DOPC membrane patch with 30 Å of water above and below and 0.05 mM KCl in solution. The final systems had ∼185 DOPC lipids, ∼96,000 water molecules, and initial dimensions of 90 x 90 x 120 Å^3^. Simulations were run using the ff14sb, Lipid17, GAFF, and TIP3P force fields in the Amber18 engine. Equilibration was carried out by lowering the restraint stepwise from 5.0 to 0.05 kcal/mol/Å^2^ over 3 ns for the protein backbone, 1 ns for sidechains, and 100 ps for lipid headgroups and heavy water atoms. Next, a voltage of −80 mV was applied to the system using eight 10 mV steps of 5 ns each. Finally, unrestrained production simulations were run for 1048 and 848 ns (wild-type and R184Q/E191Q, respectively). All equilibration simulations used the Berendsen barostat with semi-isotropic pressure coupling and a target pressure of 1 atm in the NPT ensemble, whereas voltage ramping and subsequent production simulations were run in the NVT ensemble. All simulations used a hard cutoff of 10 Å for Lennard-Jones (LJ) forces, a 2 fs timestep; long-range electrostatic interactions were treated using the Particle Mesh Ewald method.

### Proteoliposome uptake assay

VGLUT reconstitution and glutamate uptake assay were performed as previously described^20^. Briefly, purified WT and R184Q/E191Q *r*VGLUT2 were reconstituted into proteoliposomes together with a His-tagged ATP synthase holoenzyme (TF_0_F_1_) from the thermophilic *Bacillus sp. PS3* with the following protein:lipid ratios (mol/mol)> 1:40,000 for TF_0_F_1_ and 1:2000 for VGLUT2. After dialysis overnight against 150 mM KCl, 5 mM MgCl_2_, 2 mM HEPES (pH 7.3), the buffer was replaced with same fresh buffer, and dialysis continued for an additional 2 h. To remove residual detergent, the sample was incubated with 100 mg/ml Bio-Beads SM-2 resin (Bio-Rad^®^) for 2 h and used immediately for uptake assay. To measure glutamate uptake driven by a proton electrochemical gradient, 25 µl proteoliposomes containing the TF_0_F_1_ ATP synthase and 2–3.5 µg VGLUT2 were diluted into 775 µl uptake buffer (150 mM K gluconate, 5 mM HEPES-KOH pH 7.3, 3 mM Mg gluconate, 0.2 mM glutamate, 10 µCi/ml L-[3,4-^3^H]-glutamic acid [PerkinElmer^®^] and 2 mM Mg-ATP) for a final Cl^-^ concentration of 5 mM. The reactions were incubated at 30 °C for 15 mins and stopped by rapid filtration through 0.45 µm HAWP discs (Millipore^®^). The filters were washed three times with ice-cold buffer (150 mM K gluconate, 5 mM HEPES-KOH pH 7.3 and 3 mM Mg gluconate), dried, solubilized in CytoScint (MP Biomedicals^®^), and the bound radioactivity measured using a LS 6000SC scintillation counter (Beckman^®^).

### In solution digestion and mass spectrometry analysis

Aliquots (5 µl) containing 8.4 µg purified R184Q/E191Q VGLUT2 in 150 mM NaCl, 20 mM Tris, pH 7.4, 0.003% lauryl maltose neopentyl glycol (LMNG) buffer were mixed with 15 µl 0.2% Rapigest (Waters) in 100 mM triethylammonium bicarbonate buffer at pH 8. 1 µl 100 mM tris-(2-carboxyethyl) phosphine was then added to a final concentration of 4.76 mM and incubated at 56 °C for 30 minutes. This was followed by addition of 0.9 µl 500 mM iodoacetamide and 30-minute incubation at room temperature in the dark. For digestion, 1 µg of the proteolytic enzymes, either trypsin (Pierce, MS-grade, Thermo Scientific^®^), chymotrypsin (Sequencing grade, Roche^®^)) or GluC (Sequencing grade, Promega^®^), was then then added to the samples resulting in a 25 µL final digestion volume, and the samples were incubated for 12 h at 37 °C. After that, another 1 µg aliquot of enzyme was added, and the digestion continued for an additional 4 h. After digestion, the samples were acidified with formic acid to a final concentration of 10% and incubated at 37 °C for 30 min. The digests were then desalted using C18 ZipTip tips (Millipore^®^) following the manufacturer’s instructions, and eluted in 2 consecutive 7 µl aliquots of 50% acetonitrile/0.1% formic, followed by a third elution in 7 µl 0.1% formic in acetonitrile. Combined eluates were dry-evaporated and resuspended in 0.1% formic acid at 0.4 µg/µl.

For mass spectrometry analysis, samples (1 µg digest) were run in a 3 µm, 75 µm ID x 15 cm PepMap RSLC C18 EasySpray column (Thermo Scientific^®^). Acetonitrile gradients were used to separate peptides at a flow rate of 300 nl/min, as follows: 2-5% B in 3 min, 5-30% B in 62 min, 30-50% B in 5 min, 50-95% B in 5 min, where A is 0.1% formic acid in water and B is 0.1% formic acid in acetonitrile. Mass spectrometry analysis was performed using an Orbitrap Lumos Fusion (Thermo Scientific^®^) in positive ion mode. MS spectra were acquired between 375 and 1500 m/z with a resolution of 120000. For each MS spectrum, multiply charged ions over the selected threshold (2×10^4^) were selected for MSMS in cycles of 3 seconds, with an isolation window of 1.6 Th. Precursor ions were fragmented by HCD using a normalized collision energy of 30. MS/MS spectra were acquired in centroid mode with resolution 30000 from m/z=110. A dynamic exclusion window was applied which prevented the same m/z from selection for 30 seconds after its acquisition. Each sample was reanalyzed in a second LCMS injection using EThcD as the fragmentation method. In these runs we used the same parameters as described above except that the 9 most intense multiply charged ions over the selected threshold were selected for HCD (normalized collision energy as before) and then EThcD (supplemental collision energy 25).

Peak lists were generated using PAVA in-house software ^52^. All peak lists were searched against the rat subset of the SwissProt database (SwissProt.2019.07.31), using Protein Prospector ^53^ with the following parameters: enzyme specificity was set as trypsin, chymotrypsin or GluC, as appropriate for the particular sample, and up to 2 missed cleavages per peptide were allowed. Carbamidomethylation of cysteine residues, N-acetylation of the N-terminus of the protein, loss of protein N-terminal methionine, pyroglutamate formation from of peptide N-terminal glutamines, oxidation of methionine and palmitoylation of cysteine residues were allowed as variable modifications, and 3 variable modifications were allowed per peptide. Mass tolerance was 10 ppm in MS and 30 ppm in MS/MS. The false positive rate was estimated by searching the data using a concatenated database which contains the original SwissProt database, as well as a version of each original entry where the sequence has been randomized. A 1% FDR was permitted at the protein and peptide level. Spectra for peptides containing palmitoylated cysteine residues were manually inspected.

### pHluorin imaging and analysis

Dissociated hippocampal neurons from postnatal 0-1 day mice were prepared and grown as previously described ^54^. On days *in vitro* (DIV) 6, the cells were transduced with a lentivirus encoding wild-type or C60S/C62S/C64S(3CS) VGLUT2-pHluorin. At DIV 13-15, the cells were imaged using an inverted Nikon TE300 fluorescence microscope in a laminar flow perfusion and stimulation chamber. Images were collected by epifluorescence microscopy using the following bandpass filters: 470/40 nm excitation and 525/50 nm emission. Images were acquired at 1 Hz with 300 ms exposure time. At frame 15, action potentials (10 Hz for 30 s) were evoked by passing 1 ms bipolar current pulses through platinum–iridium electrodes, to yield fields of 5–10 V/cm. Cells were continuously perfused with Tyrode’s buffer (119 mM NaCl, 25 mM HEPES, 2 mM CaCl_2_, 2 mM MgCl_2_, 2.5 mM KCl, and 30 mM glucose at pH 7.4) containing 10 μM 6-cyano-7-nitroquinoxaline-2,3-dione (CNQX) and 10 μM 3-(2-carboxypiperazin-4-yl)propyl-1-phosphonic acid (APV) at 35 °C. After 140 s, total unquenched pHluorin signal was revealed by applying Tyrode’s buffer with 50 mM NH_4_Cl substituted for 50 mM NaCl.

The images were analyzed using ImageJ^55^. For each coverslip imaged, 35 synapses were manually selected using the fluorescence images obtained after NH_4_Cl application with a round region of interest (ROI) with a diameter of 7 pixels. For each time point the ROI mean fluorescence intensity was recorded and background subtracted. This data was then represented as ΔF/Δ_NH4Cl_, where ΔF is the mean fluorescence intensity for a given time point subtracting the mean fluorescence before stimulation and ΔF_NH4Cl_ is the peak fluorescence after NH_4_Cl addition, also subtracting the mean fluorescence intensity before stimulation. The normalized data was visualized using GraphPad Prism.

### Glutamate efflux

Glutamate efflux experiments were performed as previously described^39^ with a few modifications. HEK293T cells were plated into 24 well plates precoated with poly-L-lysine (0.1 mg/ml) and cells were transfected the following day with 395 ng pIRES2-EGFP pmVGLUT2, R213S pmVGLUT2 or D371N pmVGLUT2 together with 1.19 µl FUGENE-HD per well. The media was replaced one day later with fresh growth media. Glutamate flux was measured two days after transfection. First, the cells were washed twice in Ringer solution, pH 7.4 (145 mM NaCl, 10 mM glucose, 10 mM HEPES – pH 7.4, 4 mM KCl, 2 mM CaCl_2_, 1 mM MgCl_2_) before loading with 50 μM ^3^H-glutamate (2 µCi) in 200 µl Ringer’s solution for 10 minutes at 34 °C. The cells were then washed three times in ice-cold Ringer’s solution with 0.5 mM aspartate. Glutamate efflux was monitored in 200 µl warm Ringer’s solution or Ringer’s solution at pH 4.5-8.0 (145 mM NaCl, 10 mM glucose, 4 mM KCl, 2 mM CaCl_2_, 1 mM MgCl_2_ with 10 mM HEPES for pH 7.0-8.0 or 10 mM MES for pH 4.5-6.5) with 0.5 mM aspartate for 5 minutes at 34 °C. 150 µl of the medium was collected and the radioactivity measured by scintillation counting. Residual Ringer solution was aspirated, the cells lysed by the addition of 400 µl 1% SDS and the radioactivity measured by scintillation counting. The % glutamate efflux was calculated as follows:

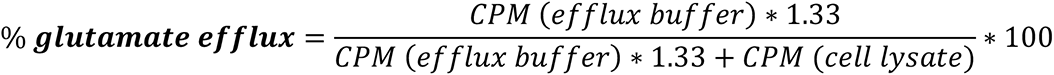

where *CPM* is the ^3^H counts per minute for the indicated samples. The data were plotted and analyzed in GraphPad Prism. Each measurement was made in duplicate and experiments performed three times independently.

### Oocyte recordings

Recordings were performed as previously described^20^. Briefly, *Xenopus laevis* oocytes were obtained from Ecocyte, injected with 50 ng pmVGLUT2-HA, R213S pmVGLUT2-HA or D371N pmVGLUT2-HA cRNA and incubated in ND96 (96 mM NaCl, 2 mM KCl, 1.8 mM CaCl_2_, 1 mM MgCl_2_, 5 mM HEPES [pH 7.4]) with 50 μg/ml tetracycline and gentamicin at 16 °C for 5 days. The oocyte currents were recorded by standard two-electrode voltage clamp in Ca^2+^ free ND96 and ND96, pH 5.5 (96 mM NaCl, 2 mM KCl, 1 mM MgCl_2_, 5 mM MES [pH 5.5]). Steady-state current/voltage (I-V) relations were obtained using a protocol of 300-ms voltage steps from −120 to +60 in 10 mV steps from a holding potential of −30 mV. Data was analyzed using Clampfit and GraphPad Prism.

### Data Availability

The atomic coordinates of *r*VGLUT2 have been deposited in the Protein Data Bank with the accession codes of 7T3N for the R184Q/E191Q structure and 7T3O for the WT structure. The corresponding maps have been deposited in the Electron Microscopy Data Bank with the accession code EMD-25665 and EMD-25665 respectively.

## Supplemental Data

**Extended Data Figure 1.**
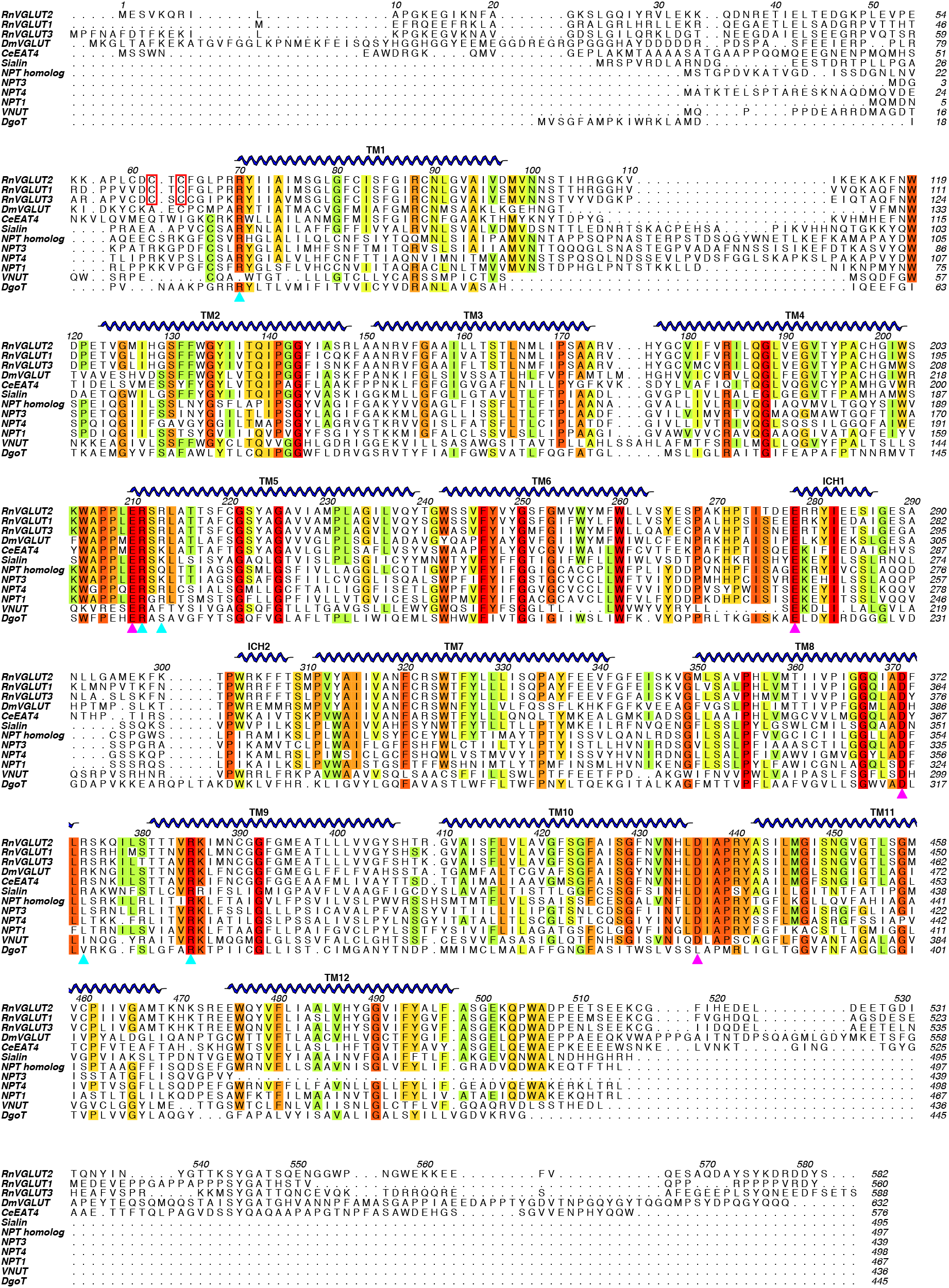
Sequence alignment of VGLUT homologs and human SLC17 family members. Sequence alignment of selected VGLUT homologs (line 1-5), other human SLC17 family members (line 6-11) and DgoT from *E. coli* demonstrates conservation across evolution and the SLC17 family of organic anion transporters. The sequences are listed in order of their sequence identity to rat VGLUT2, as in Fig. 4a. Residues are colored based on their sequence conservation, with red the most conserved and no color the least. This figure was generated with ALINE^56^.

**Extended Data Figure 2.**
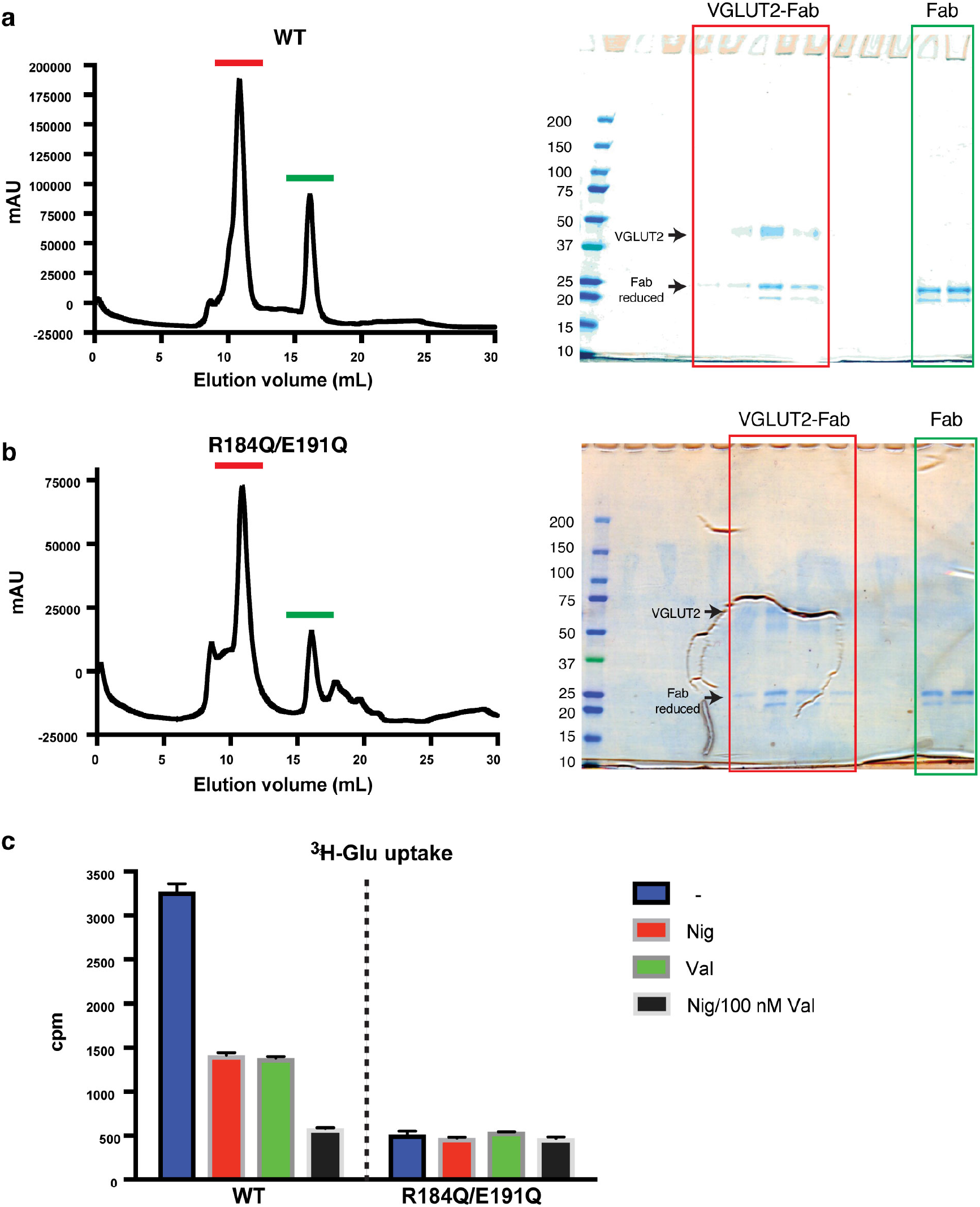
Biochemical characterization of VGLUT2-WT and R184Q/E191Q mutant. **a, b**, Size exclusion chromatography (SEC) and SDS-PAGE of WT (**a**) and R184Q/E191Q VGLUT2 (**b**) in complex with an Fab show high purity and homogeneity. Green indicates Fab alone, red VGLUT2 with Fab. **c**, Glutamate uptake by purified VGLUT2 reconstituted into liposomes with TF_0_F_1_ ATP synthase. The R184Q/E191Q mutant shows significantly less glutamate uptake than WT. The activity was measured as ATP-driven ^3^H-glutamate uptake. The ionophores nigericin and valinomycin (100 nM each) were used to dissipate ΔpH and ΔΨ, respectively.

**Extended Data Figure 3.**
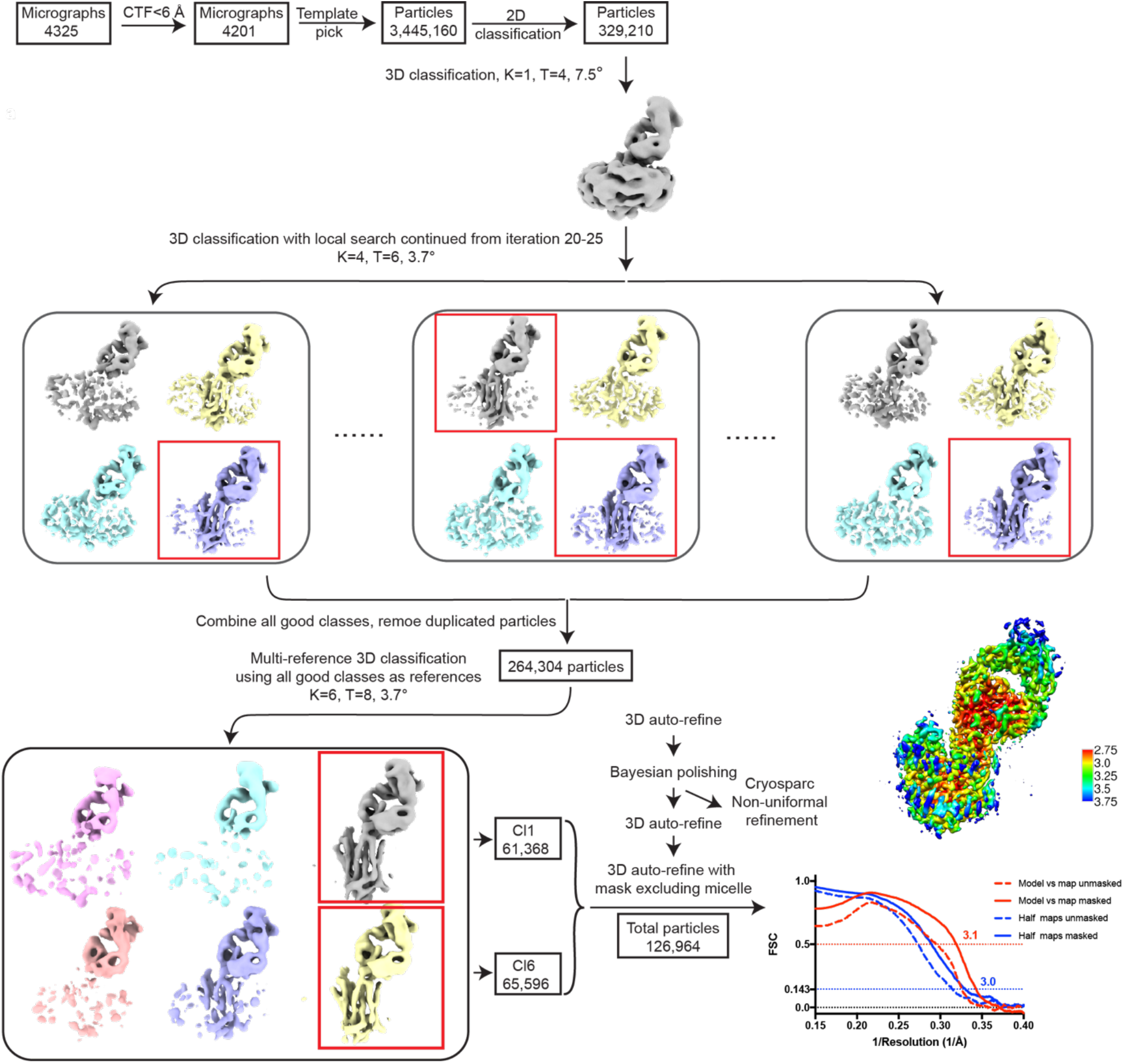
Structure determination of VGLUT2-R184Q/E191Q-Fab complex. Details of the data processing strategy are described in the Methods section. Briefly, 3 rounds of 3D classifications were carried out in RELION to select high quality particles in the same conformation. The resulting particle stack was then refined and polished in RELION. To obtain the final map, polished particles were refined using RELION 3D auto-refine with a manually generated mask including only the VGLUT2 and Fab or using the nonuniformal refinement strategy in cryoSPARC with an automatically generated mask. Both maps resulted in similar resolution and quality. Final maps were then sharpened using Phenix Auto-sharpening and used for refinement and model building. Statistics of the final map were calculated using Phenix_comprehensive_validation tool^49^ and reported in Extended Data Table 1.

**Extended Data Figure 4.**
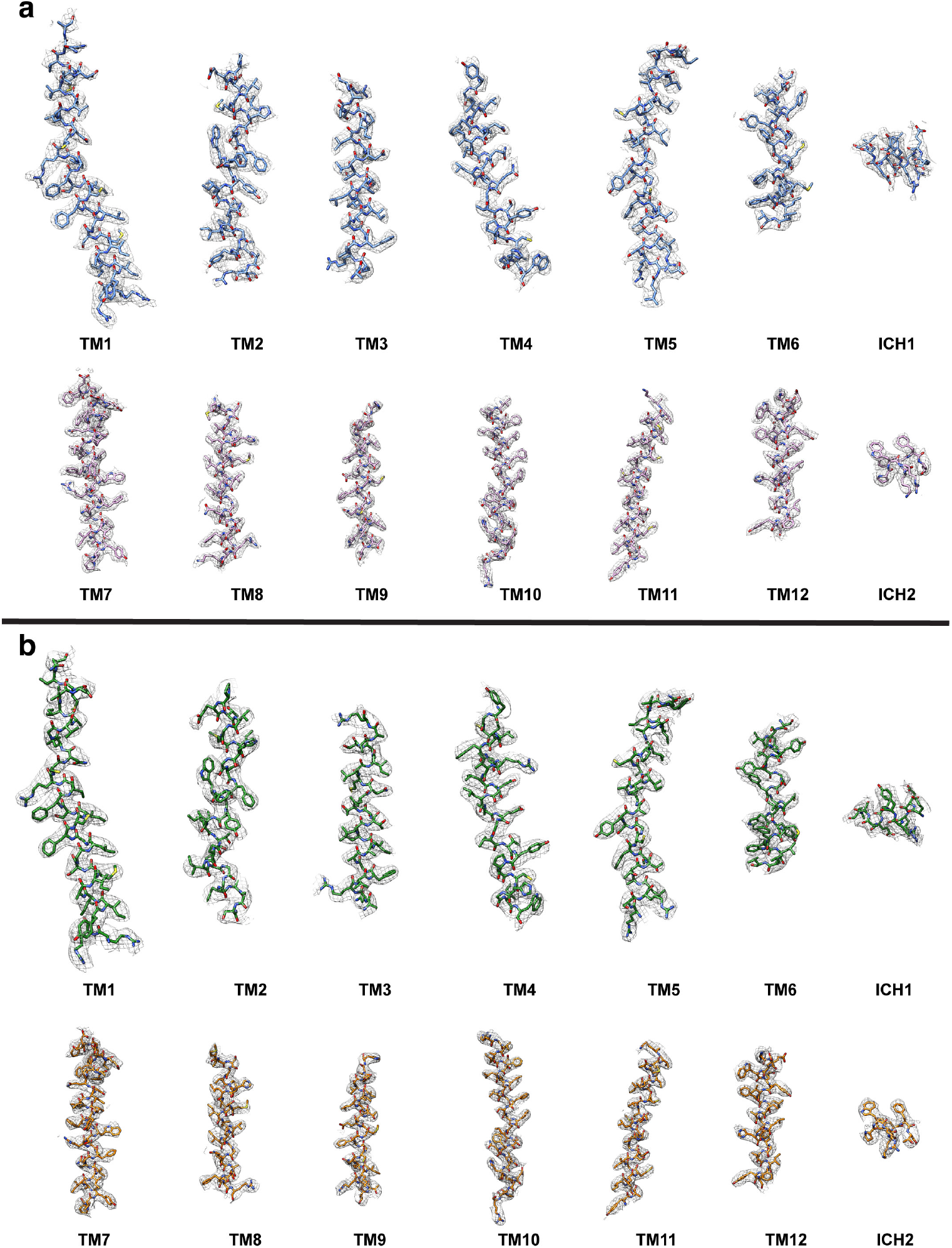
Coulombic density map for VGLUT2-R184Q/E191Q and VGLUT2-WT. Sharpened coulombic density maps of both R184Q/E191Q (**a**) and WT VGLUT2 (**b**) were of sufficient quality to resolve the side chains of most residues. The coulombic potential density maps are shown in gray mesh and models for R184Q/E191Q and WT VGLUT2 colored as in Fig. 2.

**Extended Data Figure 5.**
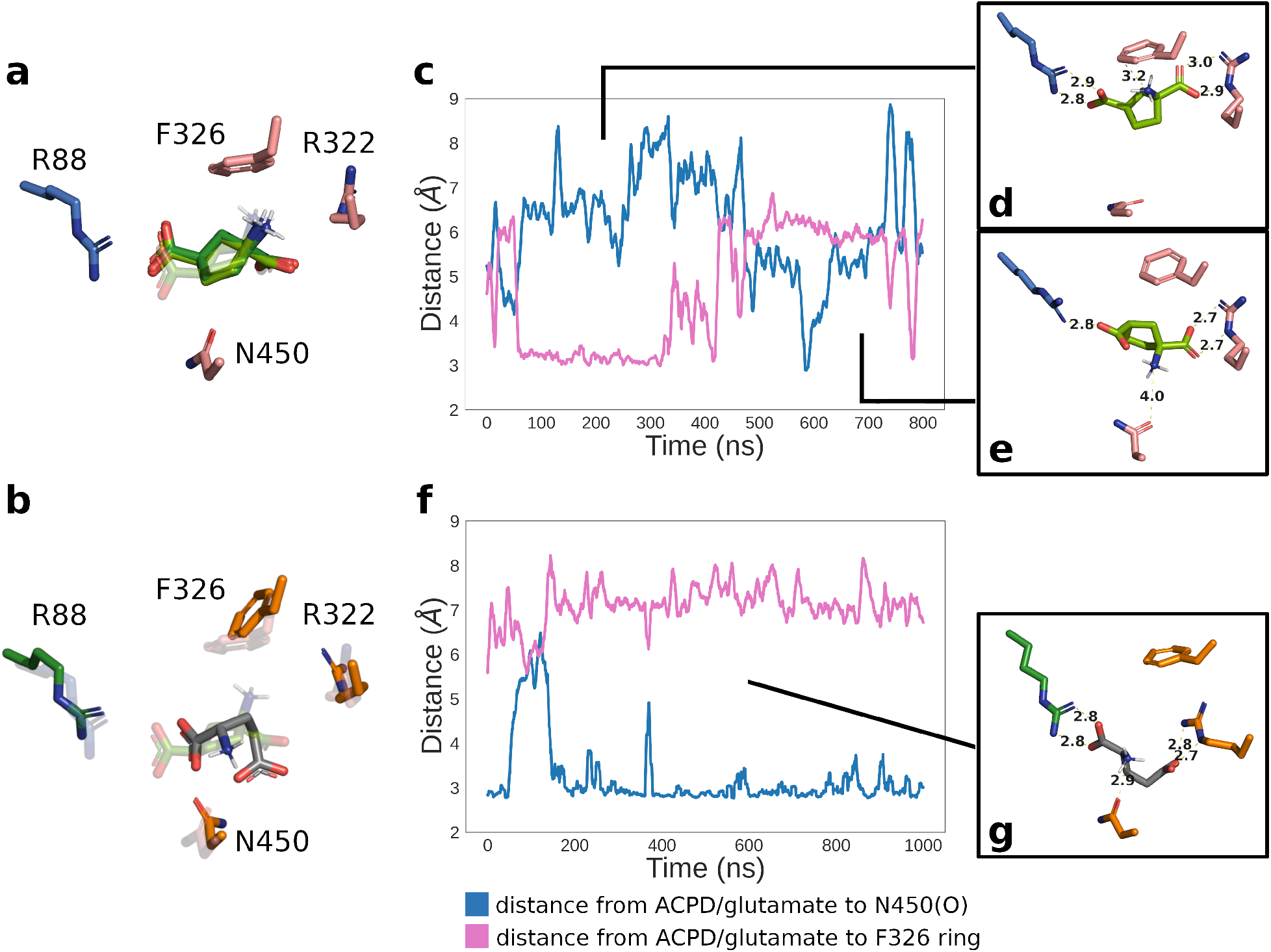
Docking and molecular dynamics (MD) simulation of ACPD and glutamate show similar interaction patterns. **a**,**b**, Docking of ACPD and glutamate. ACPD pose from the solved R184Q/E191Q-ACPD complex is shown in transparent green, interacting sidechains in blue (N domain) or pink (C domain) for R184Q/E191Q VGLUT2 (**a**), and green (N domain) or orange (C domain) for wild-type VGLUT2 (**b**). Best ACPD poses from flexible and rigid mode docking into R184Q/E191Q VGLUT2 are overlaid in opaque green and dark green, respectively. These poses are virtually identical, with all three functional groups in the same location as the solved pose (transparent). Best glutamate pose from flexible mode docking into WT VGLUT2 is shown in opaque gray (**b**). The location within the binding pocket is similar to the solved pose of ACPD (transparent green); there is an average distance of 1.5 Å between the corresponding carboxyl groups of glutamate and ACPD near R88 and 2 Å between the corresponding carboxyl groups of glutamate and ACPD near R322. (**c**, **f**), Tracking substrate recognition by MD simulation. Distance between the amine group of ACPD and the ring of F326 (pink trace) or the sidechain oxygen atom of N450 (blue trace) is shown for the 800 ns R184Q/E191Q VGLUT2-ACPD simulation (**c**) and 1000 ns WT VGLUT2-glutamate simulation (**f**). ACPD adopts two stable poses during the simulation, with the amine pointed either towards the F326 ring (**d**) or away from it and towards N450, which also removed one interaction between the carboxyl and R88 (**e**). Glutamate adopts a single stable pose during the simulation, shown in **g**.

**Extended Data Figure 6.**
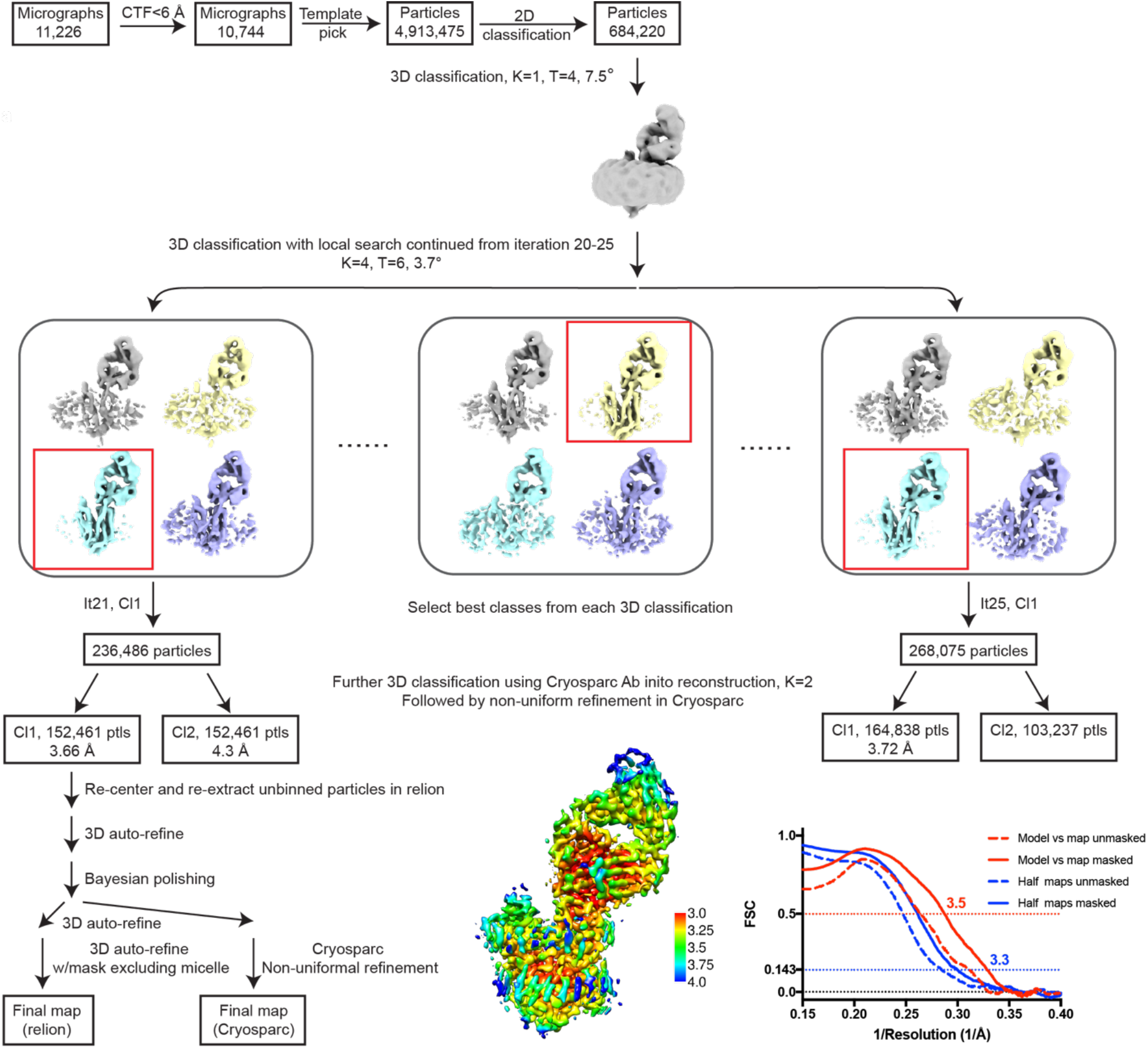
Structure determination of VGLUT2-WT-Fab complex. Details of the data processing strategy are described in the Methods section. Briefly, 3 rounds of 3D classifications were carried out in RELION and cryoSPARC to select high quality particles in the same conformation. The resulting particle stack was then refined and polished in RELION to obtain the final map. Polished particles were refined using RELION 3D auto-refine with manually generated mask including only VGLUT2 and the Fab or using the nonuniformal refinement strategy in cryoSPARC with automatically generated mask to obtain the final map. Both maps resulted in similar resolution and quality. The final map was sharpened using Phenix Auto-sharpening and used for refinement and model building. Statistics of the final map were calculated using Phenix_comprehensive_validation tool^49^ and reported in Extended Data Table 1.

**Extended Data Figure 7.**
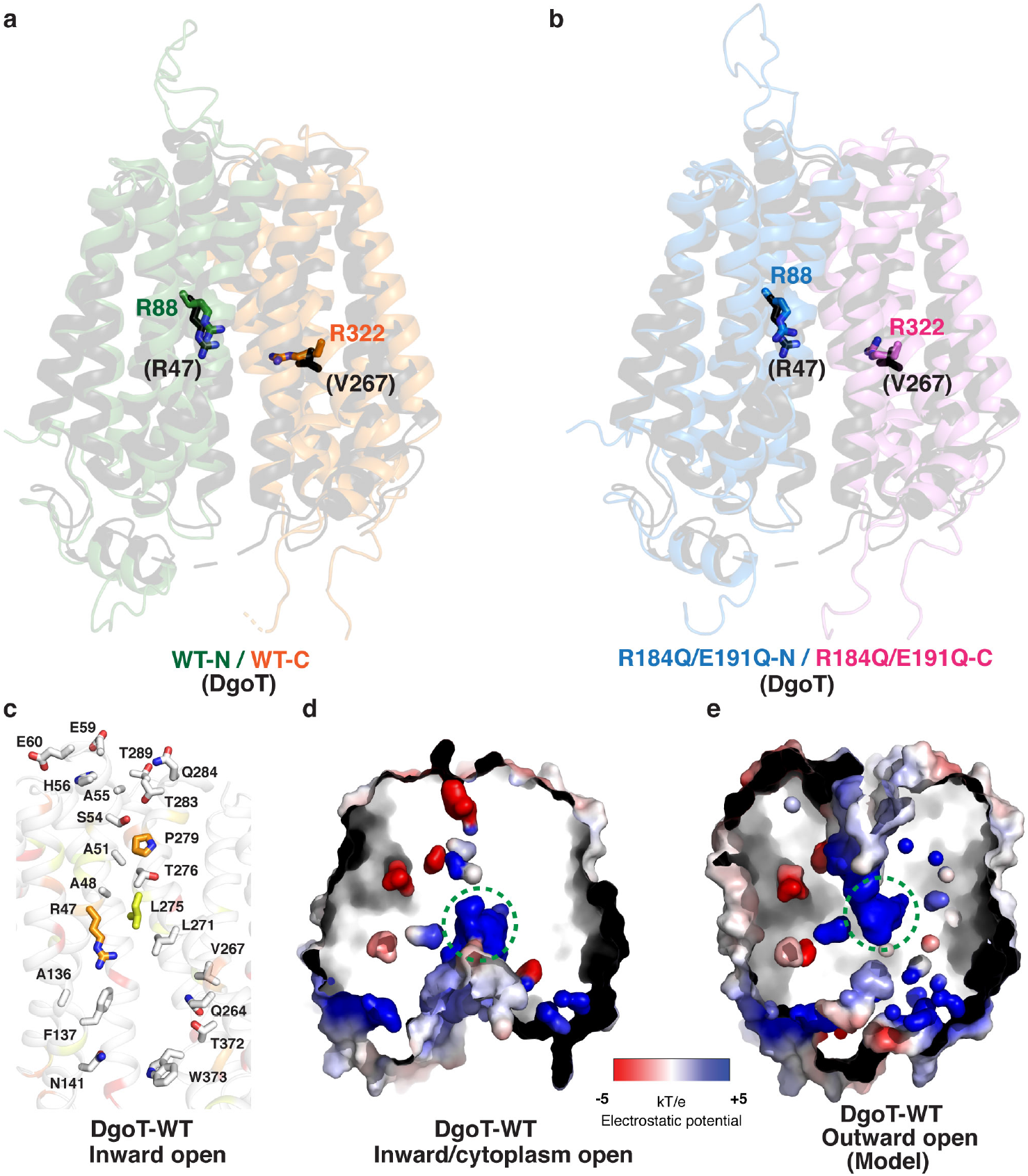
Pathways leading to the central binding cavity in DgoT are more symmetrical than those in VGLUT2. Models of cytoplasmic-open WT (**a**) and R184Q/E191Q (**b**) VGLUT2 were produced by overlapping the N and C domains separately with the crystal structure of DgoT in the cytoplasm-open conformation. The sidechains of R88 and R322 in VGLUT2 overlap well with the equivalent residues in DgoT. **c**, Structure of the binding cavity in DgoT is colored by sequence conservation (Extended Data Fig. 1), as in Fig. 3. **d**, Electrostatic surface of the inward/cytoplasm-open WT DgoT structure. **e**, Electrostatic surface of model of WT DgoT in outward-open conformation. The outward-open conformation of DgoT was generated by overlaying N and C domains separately with the structure of WT VGLUT2 in outward/lumen-open conformation.

**Extended Data Figure 8.**
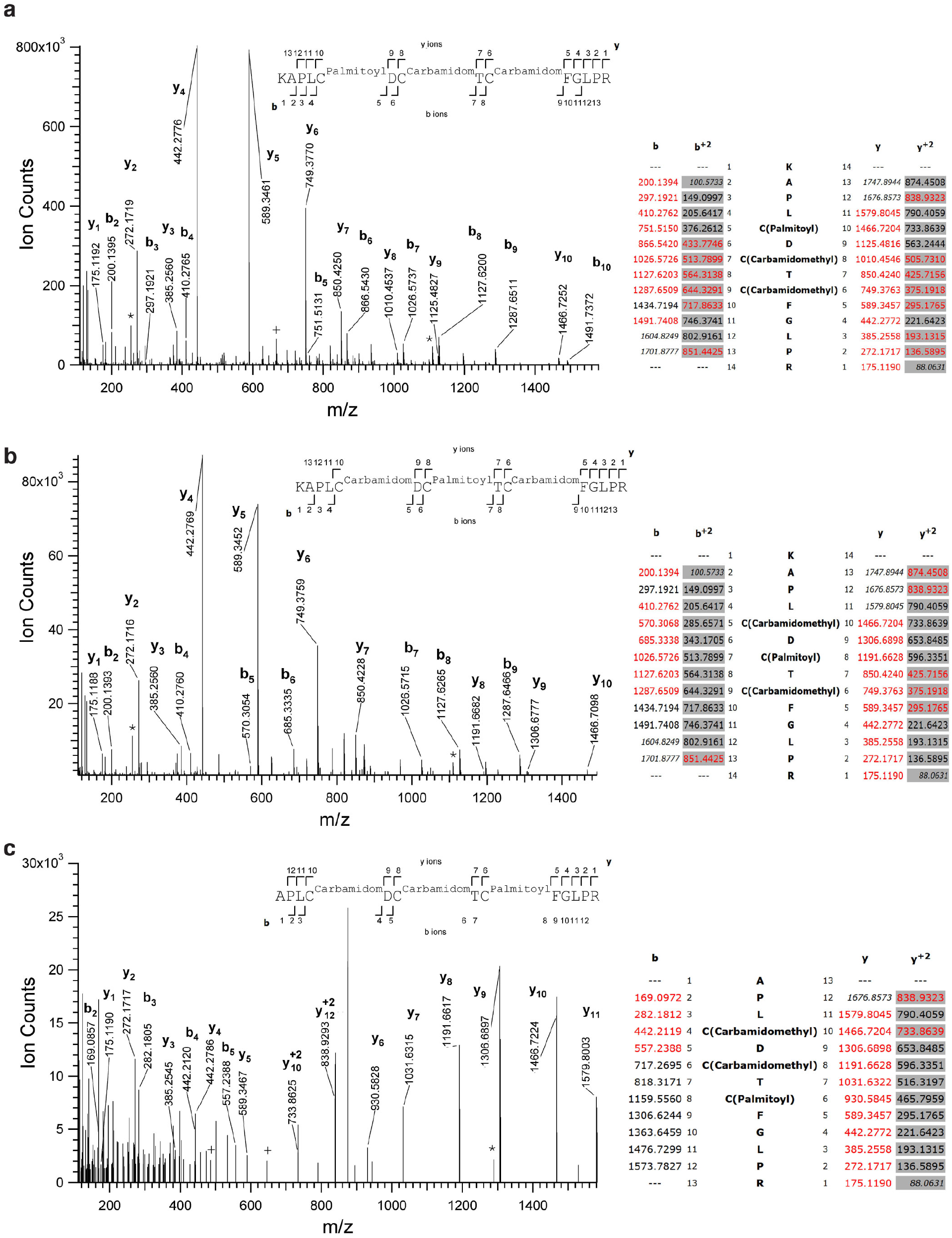
Mass spectrometry identification of palmitoylated C60, C62, and C64. MS/MS spectra of tryptic peptides spanning amino acids K56 to R69 (**a**, **b**), and A57 to R69 (**c**) of R184Q/E1891Q VGLUT2, obtained by HCD fragmentation of precursor ions 626.0019^+3^, 626.0023^+3^ and 874.4525^+2^, respectively. Excess mass of 238.229 relative to the unmodified sequence, corresponding to a palmitoyl group, was observed in the precursor and in sequence ions including particular cysteine residues. **a**, Masses of b5 [Roepstorff-Fohlmann-Biemann nomenclature] and higher and y10 and higher indicate palmitoylation at C60 in peptide K56 to R69. **b**, Masses of b7 and higher and y8 and higher pinpoint palmitoylation at C62 in peptide K56 to R69. **c**, masses observed for y6 and higher indicate palmitoylation of C64 in peptide K57 to R69. Experimental masses of the most representative sequence ion peaks are labelled in the spectra, further indicating the fragment type. The position of fragmentation events generating these ions are indicated in the sequences over the spectra. A table indicating the theoretical masses of sequence ions according to the proposed modified sequences is shown on the right, with red indicating the observed fragments.

**Extended Data Figure 9.**
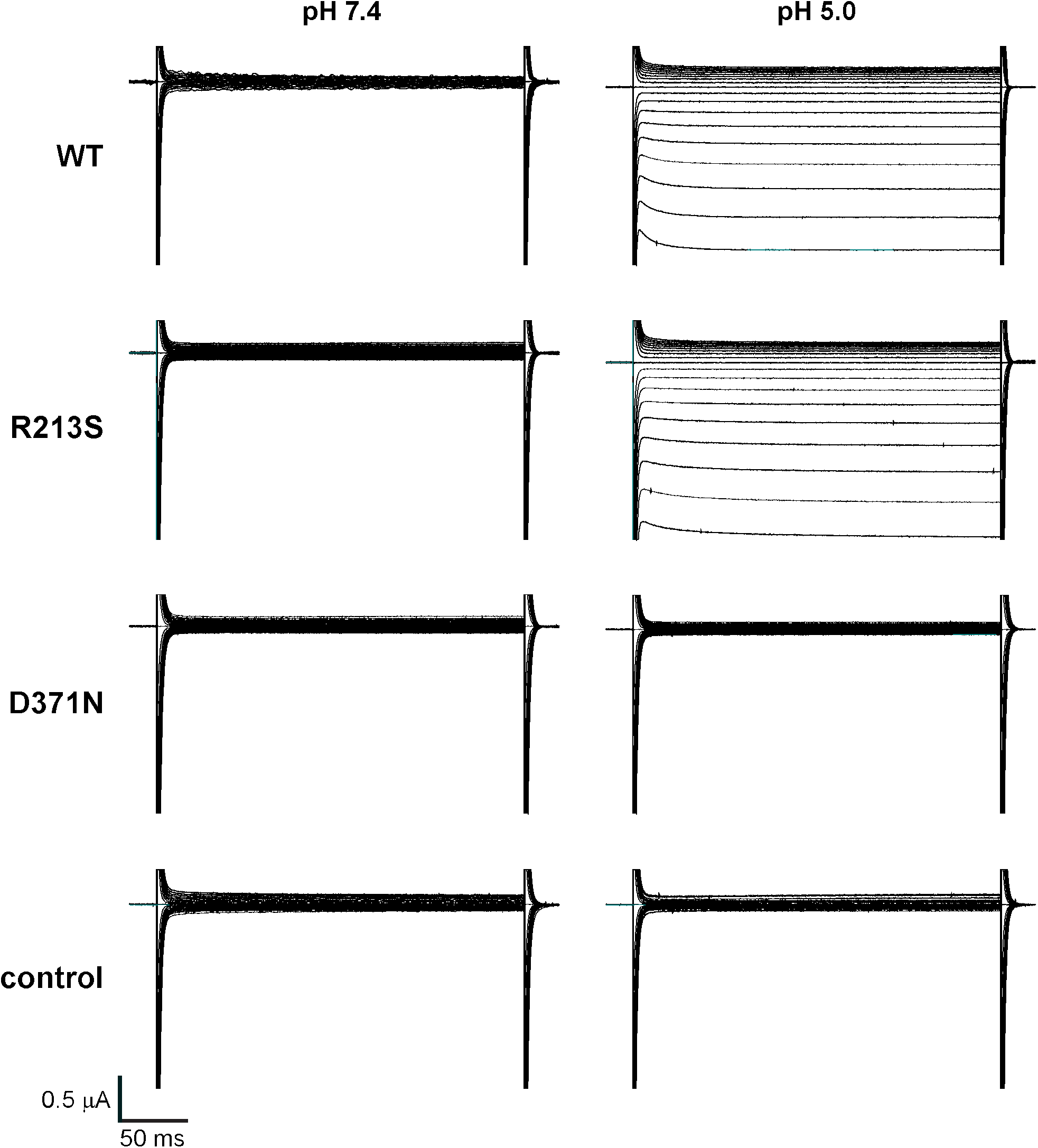
Current traces from pmVGLUT2 WT and mutant recordings. Representative current traces from oocytes injected with WT, R213S, D371N pmVGLUT2-HA cRNA and uninjected controls. Recordings were performed in steps of 10mV from −120 to 60 mV. Holding current was −30 mV.

**Extended Data Table 1.** Cryo-EM data collection, refinement and validation statistics.

|  | **R184Q/E191Q/(EMDB-25665)/(PDB 7T3N)** | **WT/(EMDB-25666)/(PDB 7T3O)** |
| --- | --- | --- |
| **Data collection and processing** |  |  |
| Microscope | Titan Krios | Titan Krios G3i |
| Camera | Gatan K3 | Gatan K3 |
| Magnification | 105,000 | 105,000 |
| Voltage (kV) | 300 | 300 |
| Electron exposure (e<sup>-</sup>/Å<sup>2</sup>) | 67 | 58.65 |
| Exposure rate | 0.57 e<sup>-</sup>/frame | 1.466 e<sup>-</sup>/frame |
| Number of frames per micrograph | 118 | 40 |
| Automation software | SerialEM | EPU |
| Underfocus range (μm) | 1.2 to 2.2 | 1.2 to 2.2 |
| Pixel size (Å) | 0.835 | 0.86 |
| Symmetry | C1 | C1 |
| Number of micrographs used | 4201 | 10,744 |
| Initial particle images (no.) | 3,445,160 | 4,913,475 |
| Refined particle images (no.) | 329,210 | 684,220 |
| Final particle images (no.) | 126,964 | 152,461 |
| Map resolution (Å) (masked/unmasked) | 3.0/3.2 | 3.3/3.5 |
| FSC threshold | 0.143 | 0.143 |
| Map resolution range (Å) | 2.75-3.75 | 3.0-4.0 |
| **Refinement** |  |  |
| Refinement package | Phenix real space refinement | Phenix real space refinement |
| Initial model used (PDB code) | 6V4D | 6V4D |
| Model resolution (Å) | 3.1 | 3.5 |
| FSC threshold | 0.5 | 0.5 |
| Model resolution range (Å) | 2.7-3.1 | 2.8-3.5 |
| Map sharpening *B* factor (Å<sup>2</sup>) | -88.06 | -62.78 |
| Model vs map CC | 0.85 | 0.85 |
| Model composition |  |  |
| Non-hydrogen atoms | 3513 | 3410 |
| Protein residues | 452 | 452 |
| Ligands | 2 |  |
| *B* factors (Å<sup>2</sup>) |  |  |
| Protein | 55.39 | 97.75 |
| Ligand | 61.07 |  |
| R.m.s. deviations |  |  |
| Bond lengths (Å) | 0.005 (0) | 0.004 (0) |
| Bond angles (°) | 0.678 (0) | 0.643 (0) |
| Validation |  |  |
| MolProbity score | 1.59 | 1.60 |
| Clashscore | 5.06 | 4.45 |
| Poor rotamers (%) | 0.00 | 0.00 |
| CaBLAM outliers (%) | 2.02 | 2.90 |
| Cβ deviation | 0 | 0.00 |
| Ramachandran plot |  |  |
| Favored (%) | 95.30 | 94.44 |
| Allowed (%) | 4.70 | 5.56 |
| Disallowed (%) | 0.00 | 0.00 |

